# Selective Nanotherapeutic Targeting of the Neutrophil Subset Mediating Inflammatory Injury

**DOI:** 10.1101/2020.06.30.180927

**Authors:** Kurt Bachmaier, Andrew Stuart, Zhigang Hong, Yoshikazu Tsukasaki, Abhalaxmi Singh, Sreeparna Chakraborty, Amitabha Mukhopadhyay, Xiaopei Gao, Mark Maienschein-Cline, Prasad Kanteti, Jalees Rehman, Asrar B. Malik

## Abstract

Inflammatory tissue injury such as acute lung injury (ALI) is a disorder that leads to respiratory failure, a major cause of morbidity and mortality worldwide. Excessive neutrophil influx is a critical pathogenic factor in the development of ALI. Here, we identify the subset of neutrophils that is responsible for ALI and lethality in polymicrobial sepsis. The pro-inflammatory neutrophil subpopulation was characterized by its unique ability to endocytose albumin nanoparticles (ANP), upregulation of pro-inflammatory cytokines and chemokines as well as the excessive production of reactive oxygen species (ROS) in models of endotoxemia and septicemia. ANP delivery of the drug piceatannol, a spleen tyrosine kinase (Syk) inhibitor, to the susceptible subset of neutrophils, prevented ALI and mortality in mice subjected to polymicrobial infection. Targeted inhibition of Syk in ANP-susceptible neutrophils had no detrimental effect on neutrophil-dependent host defense because the subset of ANP^low^ neutrophils effectively controlled polymicrobial infection. The results show that neutrophil heterogeneity can be leveraged therapeutically to prevent ALI without compromising host defense.

## Introduction

Subsets of neutrophils differ markedly in their response to both homeostatic and inflammatory signals (1). Neutrophil heterogeneity is apparent in the lungs of naïve mice, where a large proportion is marginated in the microvasculature, where they may function as immune sentinels, while other neutrophils circulate unimpeded (2). Neutrophils are an essential component of the innate immune response to polymicrobial infection due to their ability to eliminate the infectious agents (3). However, neutrophils can also become pathogenic in diseases by promoting excessive inflammation such as in the case of acute lung injury (ALI), a main cause of morbidity and mortality worldwide (4). Excessive activation of neutrophils by bloodstream bacteria and their products, such as the bacterial endotoxin lipopolysaccharide (LPS), results in tissue damage and organ dysfunction (5). Therapeutic efforts of curbing this excessive neutrophilic inflammation have been frustratingly ineffective (6). Administration of nitric oxide, norepinephrine, low dose corticosteroids, prostaglandin E1, or recombinant activated protein C, when critically evaluated, did not significantly improve patient mortality (7). Moreover, neutralizing key inflammatory mediators such as the cytokines TNF-α, and IL-1β, or reactive oxygen species (ROS), has also failed (8). In experimental models, the elimination of neutrophils markedly decreases the severity of ALI (4). On the other hand, there is the risk of compromising host defense in the setting of global neutrophil impairment and deterioration of pulmonary function during recovery from neutropenia (9).

Targeting specific subsets of neutrophils could represent an optimal therapeutic approach if one could identify deleterious neutrophil subsets without compromising the subsets essential for host defense. The unique ability of neutrophils to rapidly change their phenotype and function according to changes in their microenvironment (10-14) is a manifestation of neutrophil heterogeneity, but the distinct roles of neutrophil subsets in the setting of sepsis or endotoxemia and not well understood. We hypothesized that subsets of neutrophils are primarily responsible for the maladaptive hyper-inflammatory response that causes ALI, multiple organ failure, and death. In the present study, we identified the subset of neutrophils that incorporated specially formulated albumin nanoparticles (ANP) as the subset that could be therapeutically targeted without impairing the elimination of bacteria in experimental septicemia.

## Results

### Heterogeneous response of neutrophils to endotoxin and septicemia

After *i.v.* injection of albumin nanoparticles (ANP) to naive mice, we observed ANP-uptake in liver and spleen, whereas lungs, heart and kidney remained mostly free of ANP (Supplemental Figure 1). In response to *i.p.* challenge with the endotoxin of Gram-negative bacteria, lipopolysaccharide (LPS), uptake of *i.v*.-injected ANP in heart, kidney, liver and spleen did not increase compared to naïve mice. In lungs, however, ANP uptake increased significantly after LPS challenge (Supplemental Figure 1). Ly6G^+^ polymorphonuclear neutrophils (PMN) have the capacity to internalize ANP (15). We next determined whether uptake of ANP in the lung was restricted to Ly6G^+^ PMN and whether there was heterogeneity in the endocytosis of ANP among Ly6G^+^ PMN. In response to *i.p.* LPS, only CD45^+^ leukocytes endocytosed *i.v.* injected ANP whereas parenchymal cells (CD45^neg^) did not (Figure 1A). ANP-endocytosis was restricted to Ly6G^+^ PMN and largely absent in CD64^+^ monocytes/macrophages, NK1.1^+^ NK cells, or lymphocytes (data not shown). Pulmonary PMN endocytosed ANP in a bimodal manner, with one subset showing highly efficient uptake (ANP^high^), and the other subset demonstrating minimal to no uptake (ANP^low^) (Figure 1A). Bacterial endotoxins amplify the neutrophil activation in septicemic mice, leading to increased PMN sequestration in lungs where PMN release pro-inflammatory mediators and further enhance the recruitment of immune cells (16). Using cecal ligation and puncture (CLP), a reproducible and clinically relevant mouse model of polymicrobial infection that causes ALI, we found that in naïve control mice after 2 sequential i.v. injections of ANP only ∼4% of lung PMN endocytosed ANP as evidenced by ANP-specific fluorescence (Figure 1B). At 6h after a sham operation, laparotomy plus cecal ligation without puncture of the cecum, and 2 sequential i.v. injections of ANP, ANP^high^ PMN increased to only ∼11% (Figure 1B).Induction of severe polymicrobial sepsis by CLP, however, increased the frequency of ANP^high^ lung cells *6*-fold over baseline conditions to ∼24% (Figure 1B). We consistently found that CD11b expression levels on PMN in peripheral blood, lung, and liver, were greater on ANP^high^ PMN than on ANP^low^ PMN (Figure 1E), and lung ANP^high^ PMN showed the highest CD11b expression (Figure 1E), indicating a higher level of inflammatory activation of the ANP^high^ PMN subset. Moreover, in septicemic mice, the percentages of ANP^high^ PMN was significantly greater than in sham controls in blood, lung, and liver (Figure 1F), consistent with an increased pro-inflammatory state as well as increased adhesiveness of the ANP^high^ PMN subset. This heterogeneity in CD11b activation on PMN suggested that susceptibility to ANP-endocytosis delineated distinct subsets of pulmonary PMN.

**Figure 1.**
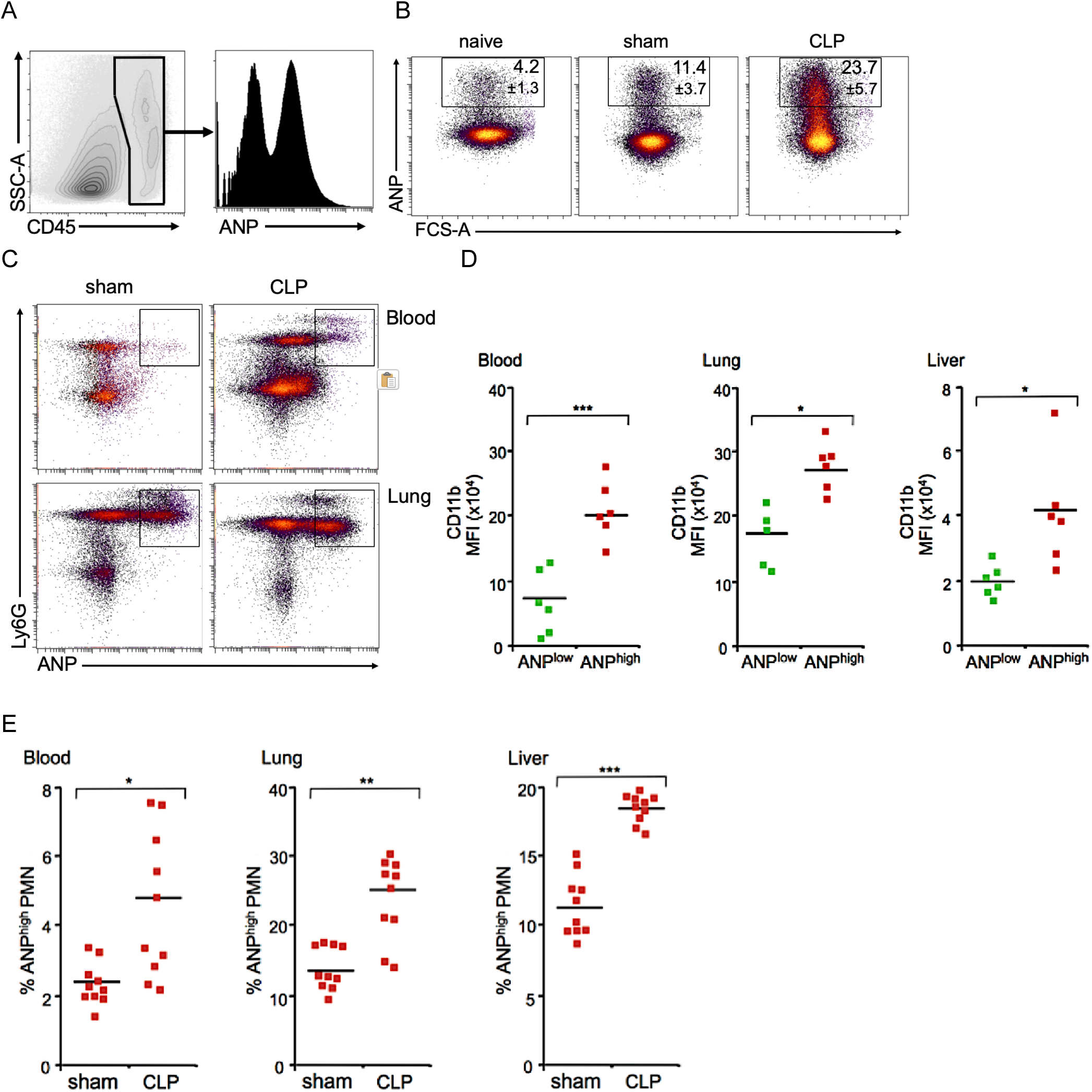
Differential susceptibility of PMN to endocytose albumin nanoparticles (ANP). **(A)** Flow cytometric analysis of lung cells from an LPS-challenged mouse. Only CD45^+^ leukocytes endocytose ANP. ANP endocytosis is bimodal, ANP^low^ or ANP^high^. **(B)** Lung cells from naive, sham-operated, or CLP-treated septicemic mice. ANP-specific fluorescence in single cell suspension of lungs. **(C)** ANP-specific fluorescence is restricted to Ly6G^+^ polymorphonuclear neutrophils (PMN) in lung and in peripheral blood. **(E)** Peripheral blood, lung, or liver cells. Mean fluorescence intensity (MFI) of CD11b-expression on ANP^high^ or ANP^low^ PMN. **(F)** Peripheral blood, lung, or liver cells. Percentage of ANP^high^ PMN is shown. Naive, sham, laparotomy plus cecal ligation without puncture of the cecum, CLP, laparotomy plus cecal ligation and puncture of the cecum. Squares represent individual mice, bars indicate mean values. * p< 0.05; ** p< 0.01; *** p< 0.001 (Student’s t test).

### ANP^high^ PMN have a distinct transcriptomic profile

To define the differences between the PMN subsets, we next analyzed whether ANP^high^ PMN had a transcriptomic profile different from ANP^low^ PMN. We performed an unbiased analysis of lung PMN transcriptomic profiles using RNA-Seq. We challenged mice with *i.p.* injections of LPS or saline and administered ANP *i.v.* 5h later. At 1h after ANP injection, we euthanized mice and harvested PMN from single cell suspension of their lungs and sorted the Ly6G^+^ PMN by flow cytometry according to their ANP uptake into ANP^low^ and ANP^high^ PMN. Immediately after sorting, we prepared PMN mRNA for RNA-Seq analysis. We generated a heat map and dendrogram (Figure 2A) to represent the normalized PMN gene expression data. We found that the biological replicates clustered into 4 groups with distinct transcriptomic profiles; i.e., the mRNA profiles defined PMN from LPS-challenged or saline-injected mice, and were distinct in ANP^low^ and ANP^high^ PMN (Figure 2A). Using MetaCore Pathway analysis to identify pathways that were different between ANP^high^ PMN and ANP^low^ PMN, we found that the pathways regulating immune response and immune cell migration were significantly over-represented in the ANP^high^ PMN (Supplemental Table). Pathways containing chemokine receptors were significantly enriched in ANP^high^ PMN, consistent with the concept that PMN heterogeneity is a function of differential PMN trafficking into tissue presumably facilitated by different chemoreceptor expression. We also found that chemokine receptors were over-represented 6.5-fold (p=0.01, Fisher’s Exact test) in ANP^high^ PMN derived from LPS-challenged mice and 4-fold in ANP^high^ PMN of naïve mice (p=0.0005, Fisher’s Exact test). Moreover, chemokines were over-represented 3.9-fold (p=0.04, Fisher’s Exact test) in ANP^high^ PMN of LPS challenged mice and 3-fold in ANP^high^ PMN of naïve mice (p=0.0013, Fisher’s Exact test).

**Figure 2.**
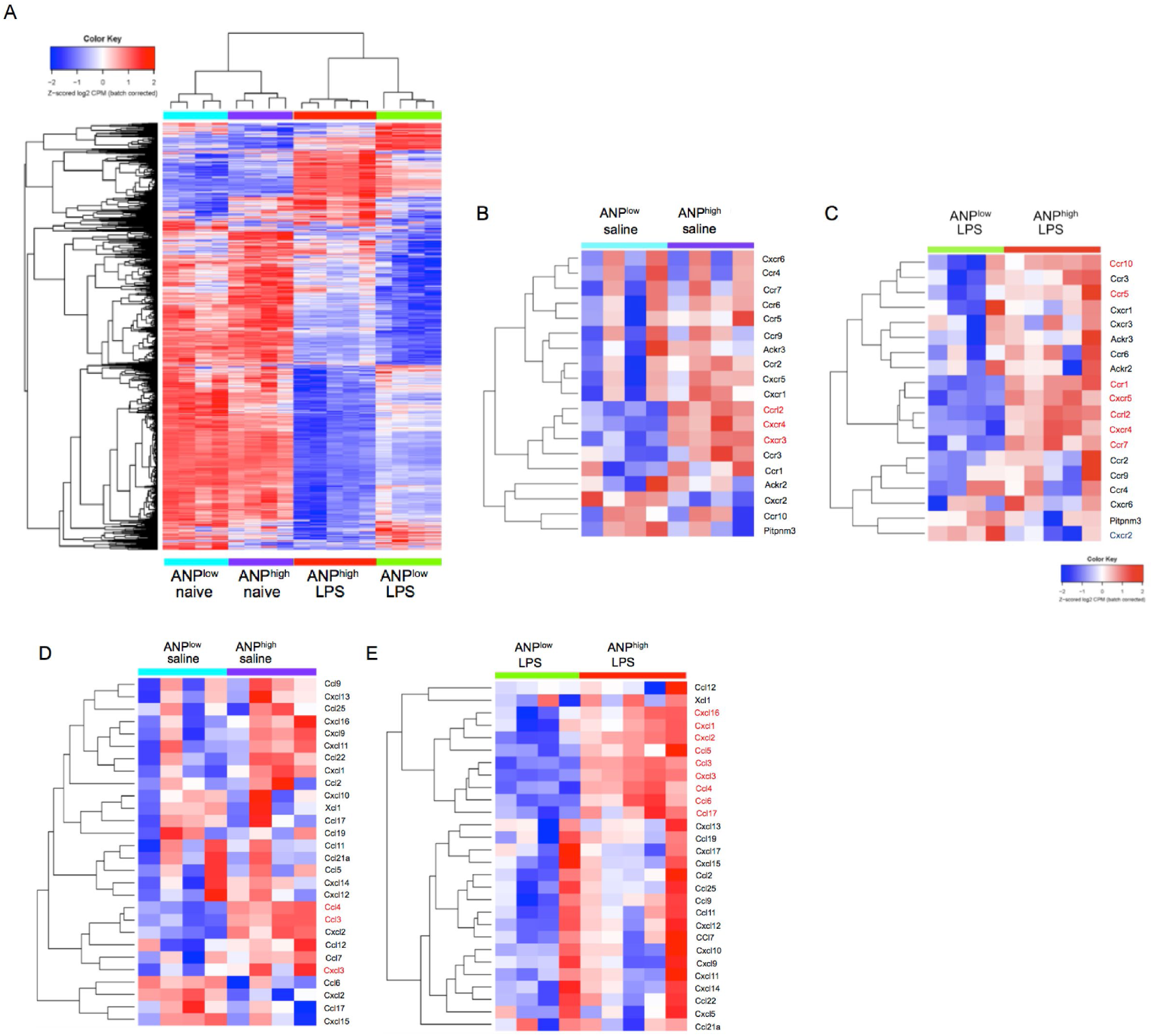
Lung PMN heterogeneity. A-E, Transcriptomic profile of lung Ly6G^+^ PMN showing high and low ANP-uptake (ANP^high^ vs. ANP^low^). **(A)** Dendrogram and heat map showing normalized gene expression data of biological replicate samples derived from lung PMN of saline injected control mice, ANP^low^ (blue), ANP^high^ (purple), or of LPS-challenged mice, ANP^high^ (red), ANP^low^ (green). **(B-E)** Heatmaps showing chemokine receptors and chemokines that demonstrate significantly higher expression levels in ANP^high^ PMN, using an FDR (False Discovery Rate) threshold of 0.05. Chemokine receptor expression in lung PMN from **(B)**, saline injected mice, or **(C)** LPS-challenged mice, or chemokine expression in lung PMN from **(D)** saline injected mice or **(E)** LPS-challenged mice. Color key, higher expression values are shown in red, lower expression values in blue. Mice were challenged for 6h with one i.p. injection of LPS (12 mg/kg) or saline; 5h after challenge, mice were injected, i.p., with one dose of fluorochrome-labeled ANP. 1h later, mice were euthanized and Ly6G^+^ PMN from lungs were sorted according to their ANP-uptake and their mRNA was processed for RNA-Seq. Representative data from 3 independent experiments.

To identify the chemokine receptors for each PMN subset, we generated separate heat-maps for chemokine receptors, plotting all genes with CPM > 0.25 (10 reads at sequencing depth of 40M reads) regardless of differential expression levels. In naïve mice, ANP^high^ PMN showed relative over-expression of chemokine receptors Cxcr3, Cxcr4, and Ccrl2 (Figure 2B). In LPS-challenged mice, ANP^high^ PMN showed relative over-expression of chemokine receptors Ccr1, Ccr5, Ccr7, Ccr10, Ccrl2, Cxcr4 and Cxcr5 (Figure 2C). We next assessed the expression of chemokines in ANP^high^ PMN and ANP^low^ PMN. In saline injected control mice, ANP^high^ PMN were significantly enriched for the expression of the chemokines Ccl3, Ccl4, and Cxcl3 (Figure 2D). In LPS-challenged mice, ANP^high^ PMN demonstrated relative over-expression of the chemokines Ccl3, Ccl4, Ccl5, Ccl6, Ccl17, Cxcl1, Cxcl2, Cxcl3, Cxcl16 (Figure 2E). Of note, mice were only exposed to ANP for the last hour of the 6h LPS-challenge, and the differences in gene expression between ANP^high^ PMN and ANP^low^ PMN in LPS challenged mice were far greater than those in the naïve mice, indicating that the ANP uptake itself likely did not affect the gene expression profiles.

These RNA-Seq data unequivocally demonstrated the existence of lung PMN subsets with a distinct response to the inflammatory stimulus LPS. Based on our RNA-Seq data and MetaCore Pathway analysis, we selected a group of chemokine receptors and cytokines to determine the kinetics of their expression after LPS-stimulation. We performed this independent validation by quantitative PCR (qPCR) and flow cytometry. We found that mRNA expression of the chemokine receptor Ccr1 was significantly greater in ANP^high^ PMN than in ANP^low^ PMN at 3h, 6h, and 12h after LPS challenge (Figure 3A). Importantly, we found that CCR1 receptor cell surface expression, consistent with the mRNA data, was significantly greater on lung ANP^high^ PMN than in ANP^low^ PMN before and 3h, 6h, and 12h after LPS stimulation (Figure 3B). CXCR2 expression was reduced at 6h after LPS-stimulation compared to 3h stimulation, CXCR4 receptor cell surface expression increased at 3h after LPS-stimulation and was greater in ANP^high^ than in ANP^low^ PMN, and decreased to expression levela of unstimulated PMN thereafter (Figure 3B). These data demonstrate that the observed mRNA expression heterogeneity translated into cell surface protein expression heterogeneity.

**Figure 3.**
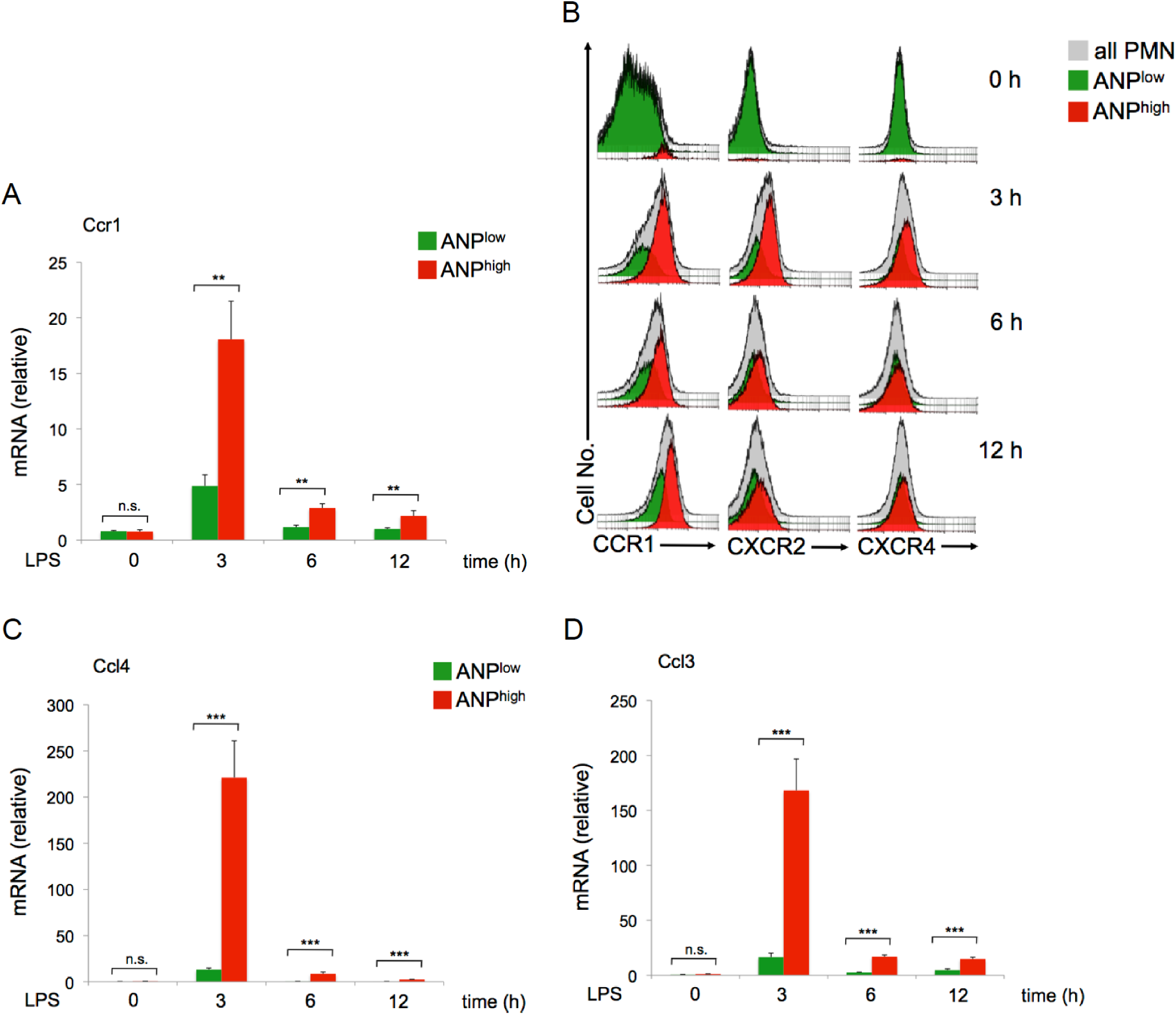
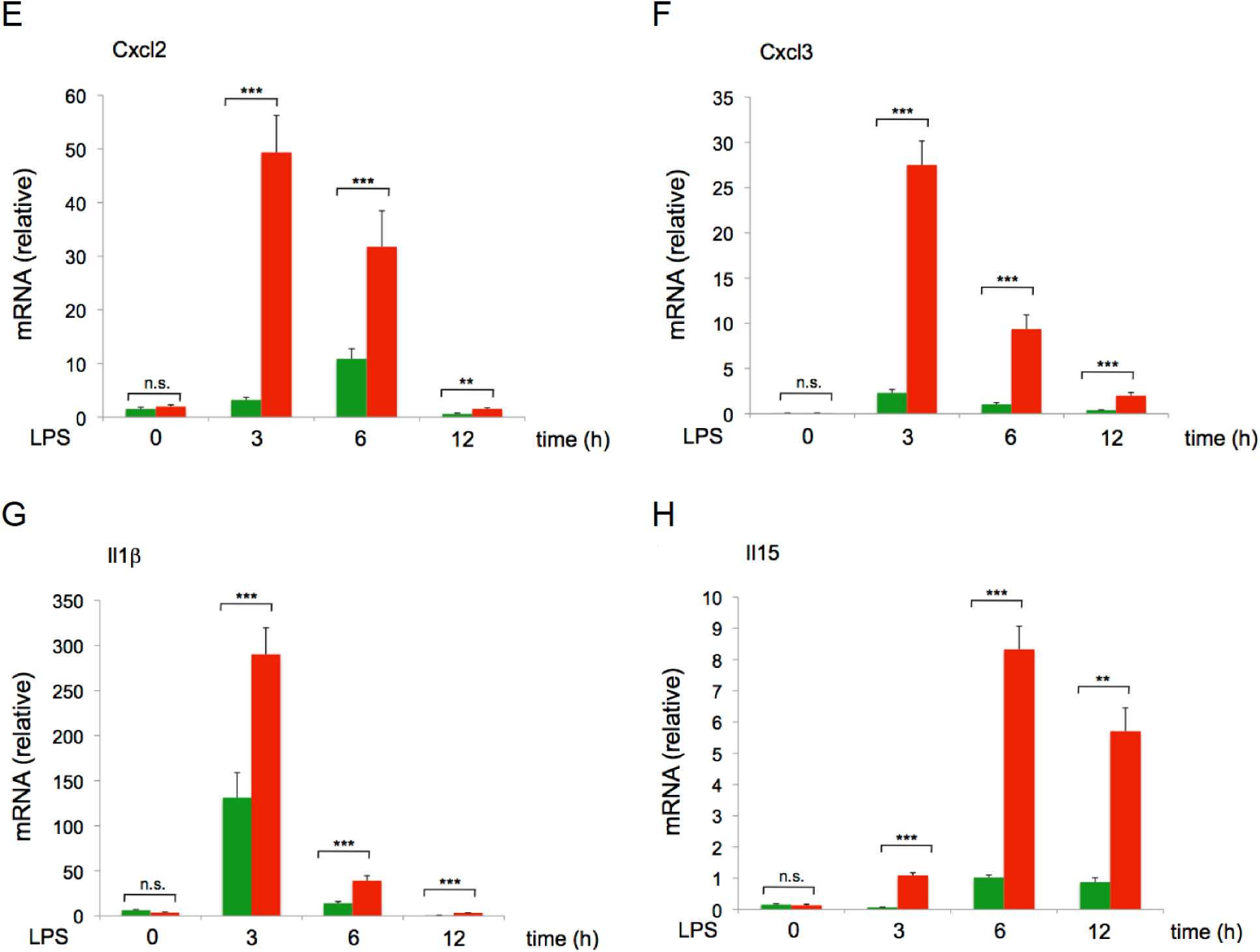
Kinetics of chemokine, chemokine receptor, and cytokine expression. Mice were treated with either saline (0h) or LPS (12 μg/ml) for 3, 6, and 12h. Fluorochrome labeled ANP were injected 1h before euthanasia. **(A)** qPCR analysis of Ccr1 in Ly6G^+^ ANP^low^ (green columns) or in Ly6G^+^ ANP^high^ (red columns) PMN. Fluorochrome labeled ANP were injected 1h before euthanasia; Ly6G^+^ PMN were sorted into ANP^low^ and in ANP^high^. **(B)** Flow cytometric analysis of lung Ly6G^+^ PMN. Lung cells were processed for flow cytometry to assess levels of cell surface expression of CCR1, CXCR2 and CXCR4 on ANP^low^ (green) versus ANP^high^ (red) PMN. Representative histograms of three independent experiments are shown. **(C-H**) qPCR analysis of selected genes in Ly6G^+^ ANP^low^ (green columns) or in Ly6G^+^ ANP^high^ (red columns) PMN. Fluorochrome labeled ANP were injected 1h before euthanasia; Ly6G^+^ PMN were sorted into ANP^low^ and in ANP^high^. (**C**,**D**) Expression of chemokines. (**C**) Ccl3; (**D**) Ccl4; Mean values and SD. n.s., not significant. ** p< 0.01; *** p< 0.001 (Student’s t test). (**E,F**) Expression of chemokines. (**E**) Cxcl2; (**F**) Cxcl3; (**G,H)** Expression of cytokines. **(G)** Il1b; **(H)** Il15. Representative data from 3 independent experiments. Mean values and SD. n.s., not significant. ** p< 0.01; *** p< 0.001 (Student’s t test)

The mRNA levels of the chemokines Ccl4, Ccl3, Cxcl2, Cxcl3 (Figure 3C-F) were significantly greater in ANP^high^ PMN than in ANP^low^ PMN 3h, 6h, and 12h after *in vivo* LPS challenge. Ccl4 (Figure 3C) and Ccl3 (Figure 3D) expression, in particular, was vastly greater in ANP^high^ than in ANP^low^ PMN, suggesting that ANP^high^ PMN are specialized cells of inflammation, which recruit and activate additional PMN. In addition, heterodimers of CCL3 and CCL4 are known to attract monocytes/macrophages (17). The cytokine IL-1β is essential for antibacterial function and expression of IL1β was induced ∼21-fold in ANP^low^ PMN 3h after LPS challenge compared to ANP^low^ PMN from saline injected control mice (Figure 3G); in ANP^high^ PMN, IL1β was induced ∼78-fold over ANP^high^ PMN from saline injected control mice (Figure 3H). Expression of the pleiotropic cytokine IL-15 expression was significantly greater in the ANP^high^ than in the ANP^low^ PMN in lungs 3h, 6h, and 12h after LPS challenge (Figure 3H). These data demonstrate that pulmonary ANP^high^ PMN can markedly amplify the inflammatory response.

### ANP^high^ PMN transfer inflammation to lungs of naive mice

We next determined whether adoptively transferring ANP^high^ PMN from donors into syngeneic recipient mice would induce inflammation in these mice. Donor BALB/c mice were challenged with a lethal dose of LPS [30mg/kg] and injected with two doses of ANP labeled with the stable fluorochrome AF647 at 1h and 2h after LPS challenge (Figure 4A). At 3h after LPS challenge, donor mice were euthanized and lung single cell suspensions were prepared for flow cytometric sorting into ANP^high^ and ANP^low^ neutrophils (Figure 4B). Syngeneic recipient mice were injected i.v. with 8×10^5^ pulmonary ANP^high^ PMN or, as controls, with an equal number of pulmonary ANP^low^ PMN from the same donors (three donors per recipient mouse were required to achieve the necessary cell number). At 2h prior to transfer of donor cells, recipient mice were treated with a sublethal dose of LPS [1mg/kg] to activate their endothelium, a prerequisite for initiating neutrophilic lung inflammation (18). At 20h after the adoptive transfer, we assessed lung inflammation in recipient mice (Figure 4A). We found ANP^+^Ly6G^+^ PMN in lungs of recipient mice, confirming successful transfer of donor cells to recipient mice and their homing to the lung (Figure 4C).

**Figure 4.**
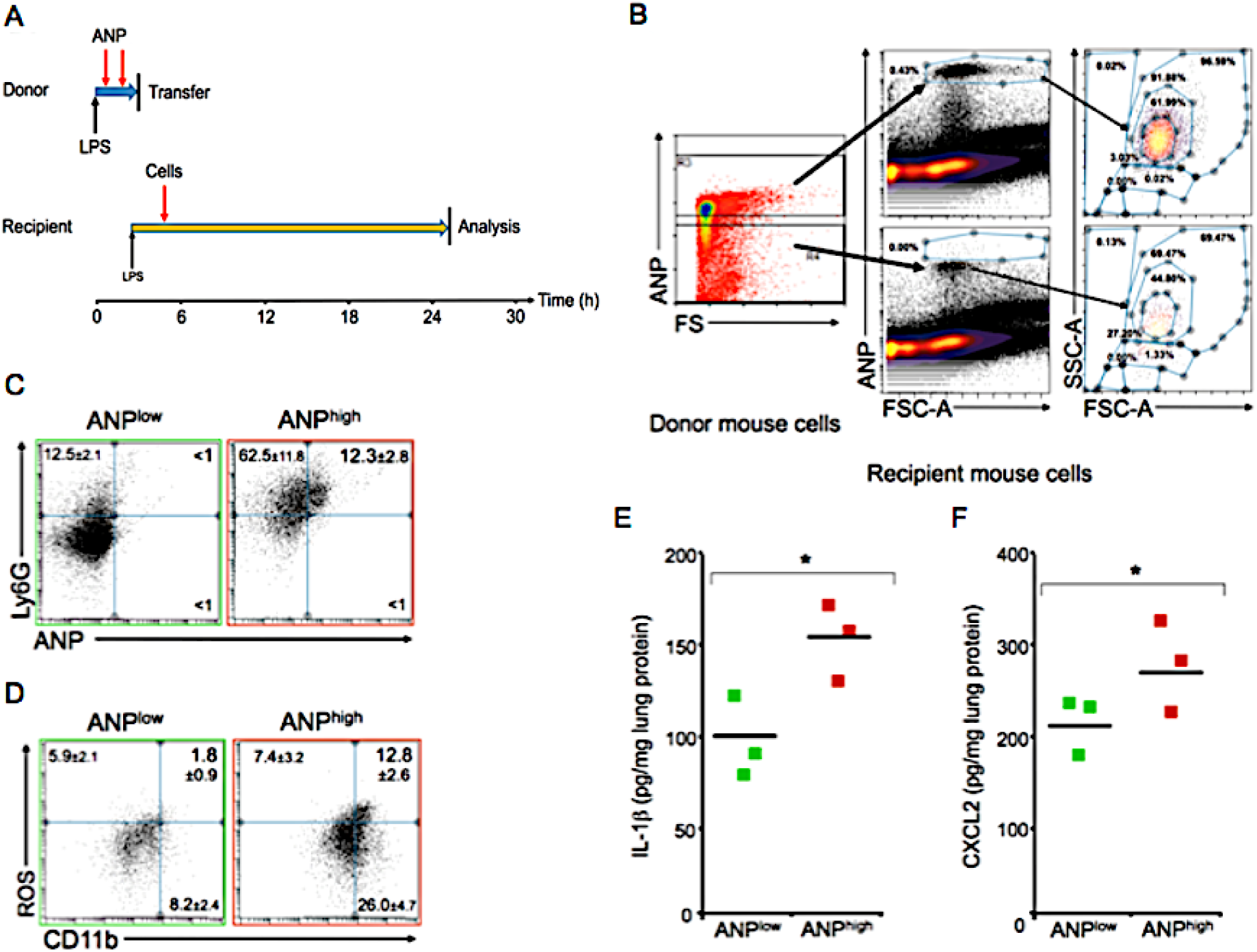
ANP^high^ PMN transfer induces inflammation in naïve lungs. **(A)** Donor mice were challenged with a lethal dose of LPS and injected with two doses of ANP labeled with the stable fluorochrome AF647, 1 and 2h after LPS challenge. 3h after the LPS challenge, donor mice were sacrificed and lung single cell suspensions enriched for leukocytes were prepared. Syngeneic recipient mice were pretreated with low-dose LPS 2h prior to adoptive transfer by intravenous injection of 8×10^5^ ANP^high^ PMN or, as controls, an equal number of ANP^low^ cells. 20h after the adoptive transfer recipient mice were sacrificed for analysis. **(B)** Flow cytometric sorting into ANP^high^ and ANP^low^ donor lung cells and flow cytometric analysis of recipient mouse lung cells. Granulocytes were gated by SSC-A^high^ FSC-A^high^. **(C)** Flow cytometric analysis of lung cells from mice that have received ANP^low^, green, or ANP^high^, red, donor cells. Dot blot showing the percentages of Ly6G ANP double- or single-positive PMN. Percentages of Ly6G^+^ANP^+^ cells were significantly greater in mice that received ANP^high^ donor cells compared to mice that received ANP^low^ donor cells: p< 0.001 (Student’s t test). **(D)** Flow cytometric analysis of lung cells from mice that have received ANP^low^, green, or ANP^high^, red, donor cells. Dot blot showing the percentages of ROS CD11b double- or single-positive PMN. Percentages of ROS^+^CD11b^+^ were significantly greater in mice that received ANP^high^ donor cells compared to mice that received ANP^low^ donor cells. **(E)** Concentrations of the IL-1β in lung tissue extracts from mice that have received ANP^low^ or ANP^high^ donor cells. **(F)** Concentrations of CXCL2 in lung tissue extracts from mice that have received ANP^low^ or ANP^high^ donor cells. p< 0.01 (Student’s t test). Representative data from 3 independent experiments using cohorts of at least 3 recipient and 9 donor mice are shown.

Transfer of donor ANP^high^ PMN significantly increased lung inflammation in the recipient mice when compared to mice that received ANP^low^ cells (Figure 4C). Moreover, after transfer of ANP^high^ PMN, lung Ly6G^+^ PMN produced more ROS when compared to controls receiving ANP^low^ PMN (Figure 3D). Because ROS induce tissue inflammation (12, 19), and ANP^high^ cells are carriers of large amounts of mRNA for inflammatory cytokines and chemokines (Figure 3), we measured the inflammatory mediators IL-1β and CXCL2 in lung tissue extracts. IL-1β, released during activation of NLRP3 inflammasome, mediates tissue injury, whereas the chemokine CXCL2 amplifies the inflammatory cycle by attracting additional pro-inflammatory neutrophils (20-22). Mice receiving ANP^high^ PMN had significantly greater concentrations of IL-1β (Figure 4E) and CXCL2 (Figure 4F) in their lungs than recipients of ANP^low^ PMN. These data demonstrated the intrinsic ability of ANP^high^ PMN to promote lung inflammation.

### Therapeutic administration of ANP^high^ PMN loaded with piceatannol protects mice from lethal polymicrobial sepsis

Integrin signaling is a main determinant of PMN behavior *in vivo.* Firm PMN adhesion on microvascular endothelium, induced by endotoxin or bacteremia, up-regulates Mac-1 (a heterodimer of CD11b and β_2_-integrin CD18), contributing to maximal activation of PMN (23). Syk activity is required for β_2_-integrin-mediated neutrophil activation (24). Given our data above, we reasoned that inhibiting integrin signaling specifically in the subset of ANP^high^ PMN would reduce lung inflammation in the polymicrobial sepsis model. We thus used the drug piceatannol, a Syk inhibitor (25, 26), that is readily incorporated into ANP due to its poor water solubility (15, 27), to inhibit Syk-mediated β_2_-integrin-dependent neutrophil adhesion. We found that therapeutic administration of piceatannol loaded ANP^high^ PMN protected CD1 mice from lethal polymicrobial sepsis (Figure 5A). Treatment with two i.v. injections of piceatannol incorporated into ANP (PANP), 2h and 4h after CLP, significantly reduced mortality of mice when compared to control groups treated with ANP without any drug after challenge with CLP (Figure 5A). Treatment using PANP reduced CLP lethality to the rate of sham-operated (laparotomy plus cecal ligation without puncture of the cecum) mice (Figure 5A). Injections of PANP alone had no effect on the survival rate compared to saline injected controls (Figure 5A). CLP challenged mice treated with ANP, without any drug, had the same mortality rate as saline-injected controls (Figure 5A), demonstrating that targeting specifically the ANP^high^ subset of PMN is sufficient to prevent CLP-induced mortality. Similarly, in the absence of polymicrobial infection but after i.p. challenge with a lethal dose of the endotoxin LPS (LD_80_), mice treated with 2 sequential i.v. injections of PANP 1h and 2h after LPS challenge, showed significantly reduced mortality when compared to control mice injected with ANP alone (Figure 5B). Reduced mortality after PANP treatment was correlated with the presence of significantly fewer highly inflammatory CD11b^high^CD45^high^ PMN (Figure 5C), and reduced CD11b expression on lung PMN when compared to ANP-treated controls (Figure 5D). Furthermore, while CD11b expression on PMN was reduced by PANP treatment in the lungs, in peripheral blood PMN CD11b surface-expression was higher when compared to PMN from ANP-treated mice (Figure 5D).

**Figure 5.**
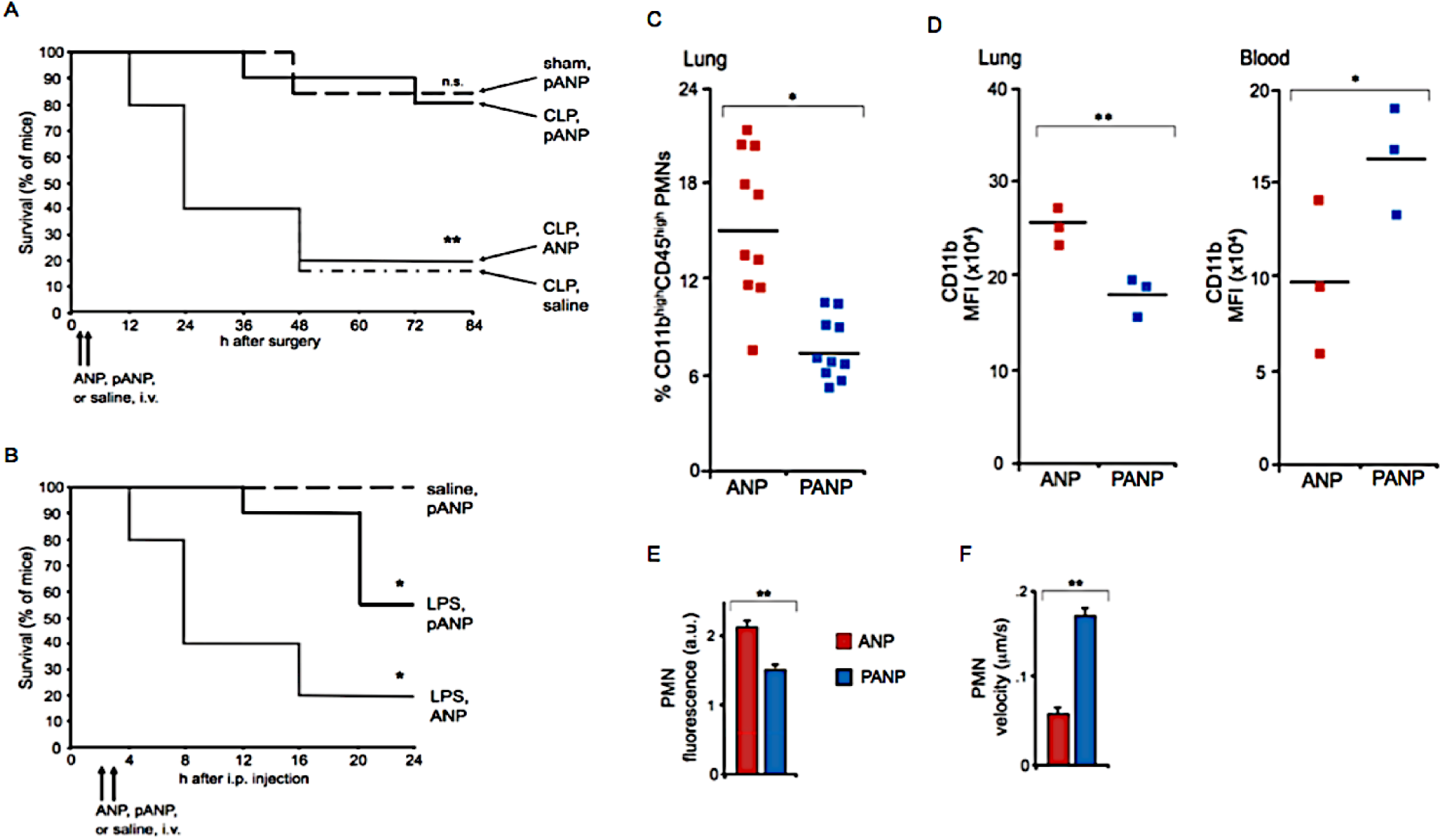
Targeting the ANP^high^ PMN prevents septicemia-induced lethality. (**A,B)** Kaplan-Meier survival curve in septicemic mice. **(A)** Polymicrobial sepsis that is lethal for 80% of saline injected or albumin nanoparticle (ANP) treated mice, is lethal in only 20% of mice after treatment with piceatannol, a Syk inhibtor, incorporated into ANP (PANP). Two consecutive i.v. injections of PANP given 2h and 4h after CLP, reduced lethality to a rate similar to sham-operated controls. **(B)** Two consecutive i.v. injections of PANP given at 1h and 2h consecutively after i.p. challenge with LPS, significantly reduced the lethality of an LD_80_ [30mg/kg] dose of LPS when compared to administration of ANP. CLP, cecal ligation and puncture; sham, cecal ligation without puncture. Representative data using a minimum of 10 mice per treatment group are shown. *p < 0.05, ** p < 0.001, n.s., not significant. (**C,D)** Flow cytometric analysis of leukocytes from septicemic mice treated with ANP or PANP. **(C)** Percentages of lung CD11b^high^CD45^high^ PMN. **(D)** Mean fluorescence intensity (MFI) of CD11b expression PMN. Leukocytes from peripheral blood or lung. (**E,F)** Two-photon microscopy of perfused, ventilated lungs of endotoxemic mice and saline-injected controls. Number and velocity of lung Ly6G^+^ PMN as a function of ANP or PANP treatment. Fluorescence, a.u., of fluorochrome-conjugated Ly6G, increases with the number of Ly6G^+^PMN. Velocity in □m/s. CD1 mice were injected with LPS [30mg/kg], 6h prior to, and with ANP or PANP 3h and 4h prior to *in vivo* imaging. **(E)** Quantification of PMN P/ANP-internalization; n=5 for ANP and n=6 for PANP; PMN number with or without P/ANP in field of view was counted and the percentage was calculated. **(F)** Quantitative analysis of PMN velocity (n=40 for ANP, n=49 for PANP). Injections of LPS and P/ANP as in (**E**). PMN velocity determined for at least 1min in the field of view. ** p< 0.01

To test whether augmented CD11b expression on PMN in peripheral blood was a consequence of PANP-induced PMN trafficking from the lung to the peripheral blood, we used two-photon microscopy to visualize PMN trafficking in the lung *in vivo* (28). We determined the number of Ly6G^+^ PMN in microvasculature and velocity of their migration through the microvasculature in lungs (Video, Figure 5E). In LPS challenged CD1 mice, the number of PMN increased, as measured by PMN-specific fluorescence (Video). Treatment of PMN with PANP, however, significantly increased the velocity of Ly6G^+^ PMN in the lung microvasculature and reduced the number of Ly6G^+^ PMN as compared to ANP treated controls (Movie, Figure 5E,F). Cell targeted treatment of ANP^high^ PMN by inhibiting β_2_-integrin signaling, and accelerated the transit of PMN through lungs, and thus reduced exposure time of lung tissue to noxious PMN-derived mediators.

### Therapeutic targeting ANP^high^ PMN improves tolerance to polymicrobial infection

Excessive ROS production is a potent mediator of tissue damage (12, 19). We found that ANP^high^ cells were characterized by high ROS production (Figure 4D). Syk, whose enzymatic activity is the cellular target of piceatannol (25), is required for integrin-mediated neutrophilic superoxide production (24). We measured ROS production by bone marrow Ly6G^+^ PMN *in vitro* (Supplemental Figure 2). Bone marrow PMN responded to stimulation with the bacterial peptide fMLP with ROS production (Supplemental Figure 2). PMN with higher uptake of PANP showed greater reduction in ROS production (Supplemental Figure 2). Moreover, the delivery of piceatannol via PANP increased drug efficacy by orders of magnitude when compared to free drug because of its incorporation in the toxic PMN subset (Supplemental Figure 2). We therefore examined whether PANP treatment reduced superoxide production by lung PMN of endotoxemic mice. We challenged mice with a lethal dose of LPS and analyzed the production of ROS by the lung Ly6G^+^ PMN *ex vivo*. We found that ANP^high^ PMN had significantly higher intracellular ROS levels than ANP^low^ PMN (Figure 6A). PANP treatment, however, almost completely blocked ROS production in these cells (Figure 6A). These data demonstrated that ANP^high^ PMN are largely responsible for ROS production by lung inflammatory cell in endotoxemia because targeted treatment via PANP markedly reduced ROS production.

**Figure 6.**
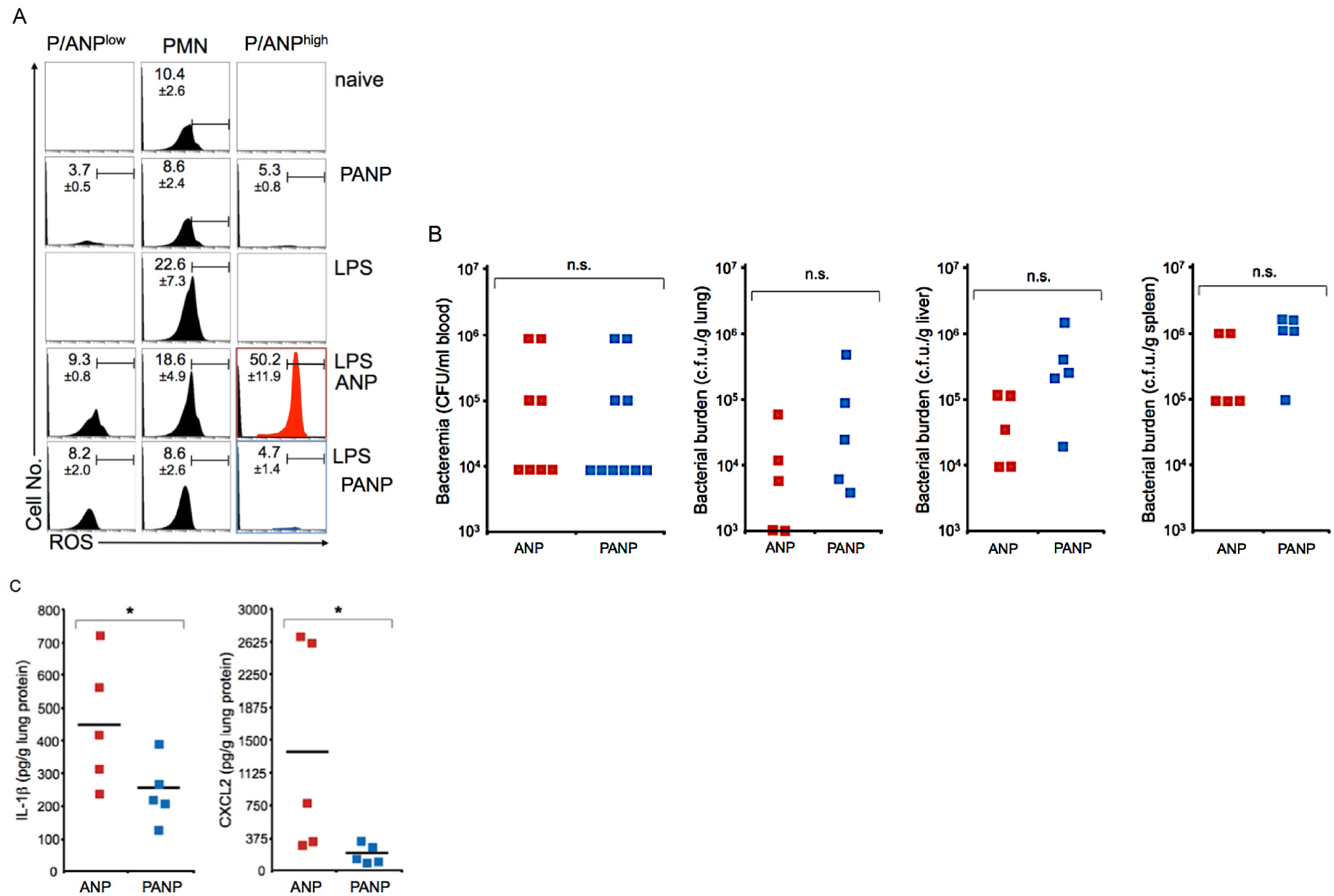
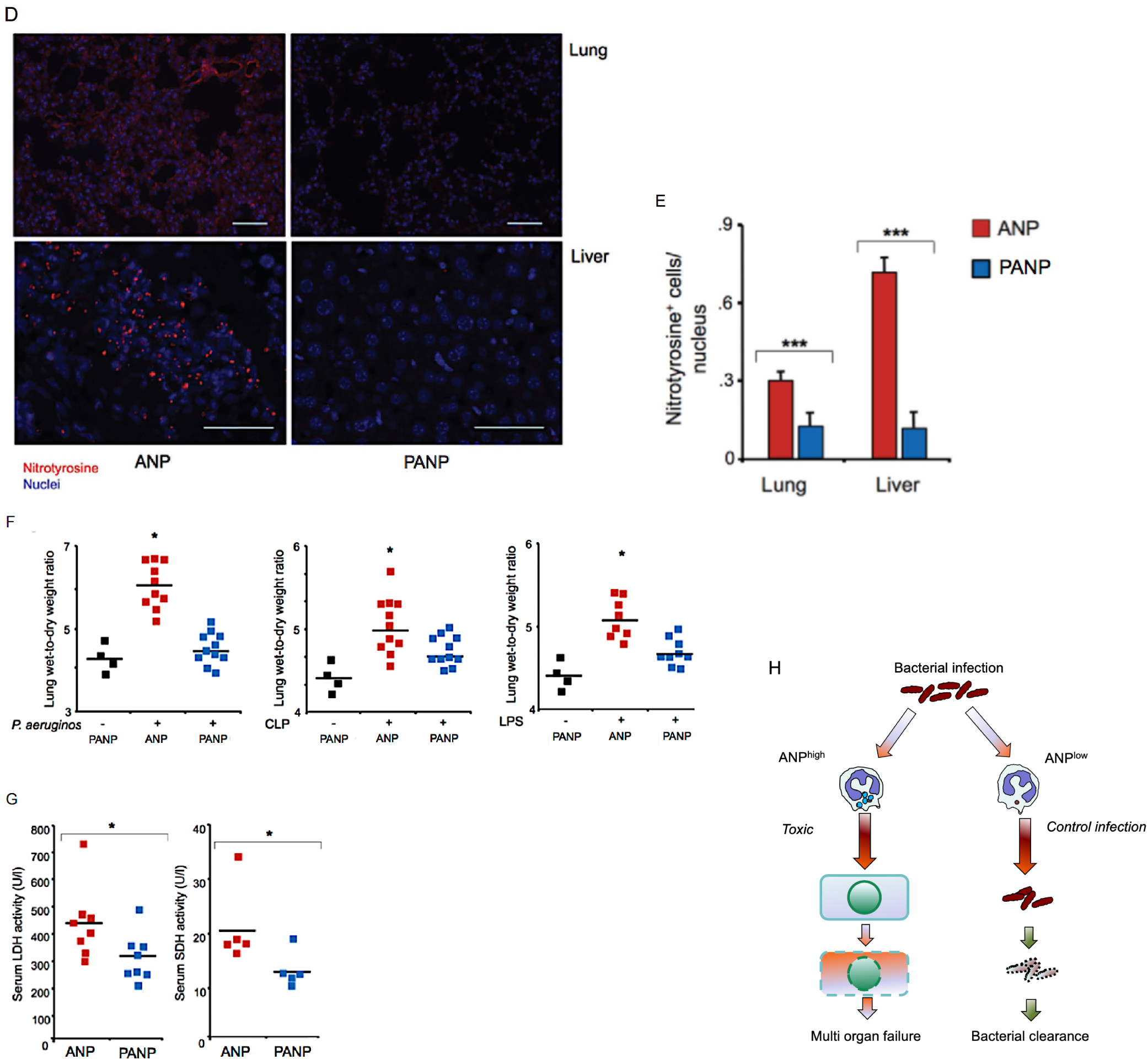
Targeting ANP^high^ PMN improves tolerance of polymicrobial infection. **(A)**Flow cytometric analysis of intracellular ROS in pulmonary PMN. Mice were treated with 2 consecutive *i.v.* injections of PANP or ANP 1h and 2h after *i.p.* challenge with LPS and ROS was measured 6h after challenge. Histograms representing ROS in all Ly6G^+^ PMN, ANP^high^ or PANP^high^ (P/ANP^high^). ROS production was measured by DHR-123 After LPS challenge, ANP^high^ PMN produce inordinate amounts of ROS (histogram shown in red), that are significantly reduced by piceatannol (PANP) treatment (histogram shown in blue). Representative data from a minimum of 3 mice per treatment group are shown. **(B)** Bacterial load (CFUs) in peripheral blood, lungs, livers, or spleens of mice post-CLP. Two consecutive *i.v.* injections of PANP given 2h and 4h after CLP did not increase bacterial burden 18h after surgery compared to ANP treated controls. **(C)** IL-1β, CXCL2 in lung protein lysate of mice 18 h after CLP. Two *i.v.* injections of PANP or ANP were given 2h and 4h after CLP. Squares represent values from individual mice and lines indicate mean values. *p<0.05. **(D)** Photomicrographs of lung or liver sections from septicemic mice treated with ANP or PANP showing nitrotyrosine formation. Paraffin embedded sections were stained with specific Ab to nitrotyrosine (red staining), and with DAPI to visualize nuclei (blue staining). Polymicrobial sepsis was induced by CLP, mice were treated with PANP or ANP 2h and 4h after challenge and were sacrificed for tissue processing and staining 18h after challenge. Bar measures 40μm, lung, or 20μm, liver. **(E)** Quantification of the presence of nitrotyrosine formation. ***p<0.001. **(F) PANP treatment reduces ALI.** Lung wet-weight to dry weight ratio of mice instilled, *i.t.*, with live *P. aeruginosa* (10^7^ CFU), or after CLP to induce polymicrobial sepsis, or after *i.p.* injections of LPS. Two consecutive *i.v.* injections of PANP, or ANP, were given 2h and 4h after challenge. Lung wet to dry weight ratio measured 6h after challenge. Squares represent values from individual mice and lines indicate mean values. *p<0.05. **(G)** Serum markers of tissue damage measured 18h after CLP and two consecutive injections of PANP or ANP given 2h and 4h after CLP. Lactate dehydrogenase (LDH) activity was significantly reduced in peripheral blood sera obtained from mice treated when compared controls treated with ANP. Hepatocyte-specific sorbitol dehydrogenase (SDH) activity was significantly reduced by PANP treatment when compared to ANP treated controls. Squares represent values from individual mice and lines indicate mean values. *p<0.05. **(H)** Model. Bacterial infection triggers two distinct activation states of PMN, characterized by endocytosis of ANP.

Mice doubly deficient for NADPH oxidase and iNOS (*gp91phox*^*−/−*^*nos2*^*−/−*^) develop spontaneous infections (29). ROS production and complementary NO production by PMN are essential to control host microbial diversity and microbial infection (29), and functional lung PMN are essential for the task of clearing bloodstream bacteria because resident macrophages in liver and spleen alone are insufficient (30, 31). We therefore determined the effects of PANP treatment on the bacterial burden of CD1 mice in the CLP model of polymicrobial infection. We found no exacerbation or amelioration of the bacterial burden as a result of PANP treatment when compared to ANP treatment (Figure 6B), suggesting that selective inhibition of integrin signaling in the ANP^high^ PMN subset did not compromise bacterial elimination. Two consecutive i.v. injections of PANP, given 2h and 4h after CLP, did not increase bacteremia when compared to ANP injected controls (Figure 6B). The bacterial burden of lungs, livers, and spleens of bacteremic mice was similar between PANP-treated mice and ANP-treated controls (Figure 6B). PMN-dependent antimicrobial function was fully preserved after PANP treatment, suggesting that antimicrobial functions, ingestion and elimination of bacteria are mainly performed by ANP^low^ PMN (Supplemental Figure 3).

Because PANP treatment did not weaken anti-microbial resistance, we analyzed parameters of tissue inflammation and damage We measured crucial inflammatory mediators, IL-1β, and CXCL2. In lung tissue extracts of mice subjected to CLP. We found a substantial reduction in the concentration of IL-1β and CXCL2 after PANP treatment when compared to ANP treated controls (Figure 6C). We next measured nitrotyrosine formation in lungs and livers of septicemic mice. In the experimental mice as well as in septicemic patients, activated lung myeloid cells, inflammatory or resident, and epithelial type II cells, release both NO and superoxide which react to form peroxynitrite, a potent oxidant causing tissue damage (32). Peroxynitrite (ONOO^*−*^), but not NO or superoxide alone, nitrates tyrosine residues (32). We observed that nitrotyrosine-specific staining in inflammatory and parenchymal cells was significantly reduced in lungs and livers of mice treated with PANP when compared to ANP-treated controls (Figure 6D,E). These data suggest that antimicrobial function and tissue damaging function are performed by distinct subsets of PMN.

We next determined the effect of PANP treatment on pulmonary edema which is a characteristic feature of inflammatory lung injury. A marked increase in lung wet-to-dry weight ratio is indicative of breakdown of the alveolar capillary barriers, the hallmark of ALI. Pneumonia is the most common cause of ALI in patients (33) and also the most common cause of sepsis (8). In the model of pneumonia induced by i.t. instillation of live *P. aeruginosa* bacteria, PANP treatment significantly reduced pulmonary edema when compared to treatment with the control ANP (Figure 6F).

Furthermore, treatment with PANP significantly reduced pulmonary edema in endotoxemic or septicemic mice when compared to lungs from ANP treated controls (Figure 6F). A reduction of tissue damage, because of reduced lung inflammation, could be the proximate cause of reduced inflammatory lung injury after treatment. Measuring a markers of overall cell damage, lactic dehydrogenase (LDH) (34), revealed that the polymicrobial sepsis-induced increased serum activity of LDH was significantly reduced by PANP treatment when compared to ANP treated controls (Figure 6G). In addition, hepatocyte-specific sorbitol dehydrogenase (SDH) activity, a marker of hepatocyte damage (35), was also significantly reduced by PANP treatment of septicemic mice (Figure 6G). Endocytosis of ANP delineates two distinct subsets of PMN. ANP^high^ PMN cause tissue damage whereas ANP^low^ PMN control microbial infection (Figure 6H).

## Discussion

Targeting neutrophilic inflammation that is pathogenic in ALI remains an important unmet clinical need. Our observation that neutrophils primed by bacterial infections or bacterial derived products exhibit heterogeneity in their capacity to endocytose ANP lead to the discovery of two distinct neutrophilic subsets, one that causes inflammatory injury and one that controls microbial infection. Functional and phenotypic PMN heterogeneity (36-40) led us to test the hypothesis that immunopathology and severe tissue inflammation of endotoxemia and septicemia can be treated by targeting a distinct neutrophilic subset. Increased expression of chemokines and chemokine receptors in ANP^high^ PMN was consistent with their role in promoting tissue inflammation. Several of the chemokines such as CCL3 and CCL4 or CXCL2 and CXCL3 are members of the macrophage inflammatory protein family and are typically thought to be released by macrophages to increase the influx of pro-inflammatory cells such as neutrophils (41). Our data suggest that a subset of neutrophils might be a substantial source of these inflammatory proteins in models of ALI. All cells need to express genes that are required for their intrinsic functions, whereas production of secreted factors can be delegated to subsets (42). Our results indicate that a specific subset of phenotypic and functionally distinct neutrophils is responsible for lethality in experimental polymicrobial sepsis. ANP^high^ PMN were characterized by higher expression of the chemokine receptors for the ligands released following LPS activation such as CCR1 and CCR5 (the receptors for CCL3) which could point to a possible positive feedback loop in which inflammatory ANP^high^ PMN attract additional inflammatory PMN, and thus actively promote a vicious cycle of hyper-inflammation and tissue injury (43). Given their phenotypic and functional profile, ANP^high^ PMN might play a pathogenic role in COVID-19, the disease caused by coronavirus SARS-Cov-2 (44). The main cause of COVID-19-mortality is acute respiratory failure (45). In patients with severe COVID-19, activated neutrophils, recruited to the pulmonary microvasculature, produce histotoxic mediators including ROS (46). Activation of neutrophils might contribute to cytokine release syndrome (“cytokine storm”) that characterizes severe COVID-19 disease (47). Therapy targeting ANP^high^ PMN might prevent a patient’s hyperinflammatory response to SARS-Cov-2 without weakening the antiviral response.

It has been shown that the incorporation of denatured albumin beads depends on Mac-1 expression (48). ANP-incorporation, by contrast, is Mac-1-independent (15) suggesting a distinct molecular mechanism of ANP-endocytosis. “Aged neutrophils”, were first described *in vitro* as functionally deficient (49) and have been subsequently shown to promote disease *in vivo* in models of sickle-cell disease or endotoxin-induced septic shock (50). While “Aged neutrophils” home to the bone marrow under steady-state conditions ANP^high^ PMN do not home to the bone marrow under steady-state conditions and markers of aged peripheral blood neutrophils, CXCR4, CXCL2, CD62L, and TLR4, do not distinguish ANP^high^ from ANP^low^ PMN under steady-state or inflammatory conditions in the lung (Figure 2). These findings thus show that the previously described subset of aged neutrophils is distinct from the ANP^high^ PMN subset we identified.

Administration of ANP carrying piceatannol, a Syk inhibitor, dramatically improved survival in polymicrobial sepsis but, critically, did not increase the host’s bacterial burden. Syk function is required for the essential antibacterial functions of neutrophils (51). We found that ANP^low^ PMN are more efficient in ingestion and elimination of *E. coli* bacteria *in vitro* than ANP^high^ PMN and inhibition of Syk in ANP^high^ PMN does not impair control of polymicrobial infection *in vivo*. By therapeutically leveraging the preferential ANP uptake of the inflammatory PMN subset, we succeeded in limiting immunopathology caused by polymicrobial infection. Pathogens adapt to host resistance mechanisms, but not host tolerance mechanisms (52). It is conceivable that an ANP-based approach of targeting toxic subset of neutrophils, by improving host tolerance, might assist the host in combating antibiotic-resistant microbial infections (53). ANP could also be used to deliver compounds other than piceatannol or to specifically deliver siRNAs or microRNAs. Furthermore, ANP-based therapy might be useful in other forms of neutrophilic injury; e.g., ischemia reperfusion injury of the myocardium or the kidney (54).

The ability to tolerate pathogens in experimental polymicrobial sepsis was greatly strengthened by targeting ANP^high^ PMN with a drug that accelerates the velocity of pulmonary PMN and abrogates their ROS production. PANP did not target β_2_-integrins directly (15) but mitigated downstream β_2_-integrin signaling, and are thus more likely to be effective when β_2_-integrin are already engaged; i.e., in the lung microvasculature of septic patients (55). Further potential improvement of ANP-based therapies over the current standard treatments of septic patients (56) lies in the precision targeting of the relevant pathogenic subset of neutrophils. Earlier efforts to neutralize reactive oxygen species, to use antibodies against key inflammatory cytokines such as TNF-α, IL-1β, or to inhibit the endotoxin LPS (8, 57-59) have failed to reduce mortality associated with ALI/ARDS possibly because they did not discriminate between PMN subsets and may have compromised both host defense (resistance) and tissue repair (tolerance) (33). Compared with the therapeutic administration of exogenous cells, generated from donors or from the patients themselves, for example, mesenchymal stromal/stem cells (MSCs) (60), ANP-based therapy has the advantage of targeting the host’s endogenous cells dependent on their pathogenic activation.

One limitation of our approach is that we cannot distinguish whether the specification into ANP^high^ PMN occurs in the bone marrow prior to egress of PMN into circulation or whether it occurs in the tissue itself. It is also possible that the ANP^high^ state characterized by upregulation of chemokine receptors and chemokine ligands is reversible and that PMN can transition between these states, in response to environmental cues even during the short life-span of neutrophils (61).

The present data support the concept that the generation of heterogeneous PMN subpopulations evolved as a host defense mechanism to avoid an indiscriminate response by all PMN to septicemia (42).A heterogeneous response of PMN to bacterial challenge below the threshold of septicemia may be sufficient to contain the infection and avoid immunopathology by striking a balance between an inflammation-amplifying PMN subset and a pathogen-eliminating PMN subset. However, excessive systemic infection may overwhelm this adaptive mechanism and shift the balance towards excessive activation of the pro-inflammatory PMN subpopulation that via the positive feedback loop of attracting even more PMN via chemokines causes tissue damage. Inhibiting Syk-dependent functions of this subset of inflammatory PMN as described would reduce damage, increase tolerance, and have no detrimental effect on the anti-microbial function of the host. The data demonstrate that sepsis represents an excessive, inappropriate activation of a subset of PMN and not an all-out indiscriminate response by PMN to septicemia. The separation of host defense function from tissue inflammatory function is reflected in the heterogeneous ability of PMN to incorporate ANP. Co-option of distinct neutrophil subpopulations by cancers and other diseases contributes to disease progression (1, 62). We demonstrate here that neutrophil heterogeneity can be effectively leveraged for a nanoparticle-based therapy.

## Materials and methods

### Mice

We used outbred male CD 1 mice, at a body weight ranging from 34g to 38g and, for adoptive transfer experiments, male inbred BALB/c mice between 8 and 10 wk of age. Mice were treated in accordance with the NIH Guide for the Care and Use of Laboratory Animals and UIC animal care committee’s regulations. All procedures were approved by the UIC IACUC.

### Synthesis of uniform-sized spheric albumin nanoparticles

ANP **and PANP**, synthesized as described (15), were of consistent **hydrodynamic size** (120nm ± 28nm diameter and **zeta potential** (−27± 5.48mV) distribution. We injected i.v. 8.3mg/kg body weight of ANP, or of ANP loaded with 8.9μM piceatannol (PANP) 1h and 2h after challenge with LPS or 2h and 4h after CLP. **Flow cytometry and cell sorting**. Single cell suspensions were prepared as described (63). Cells were stained for 30 min on ice. Dead cells were excluded by F-SC, S-SC. Neutrophils were gated by Ly6G^hi^ CD115^lo^ S-SC^hi^. Antibodies were from: Bdbiosciences, CCR1; CXCR2; CXCR4; TCR-β (H57-597); NK-1.1 (PK136); CD16/CD32 (2.4G2). ebioscience, CD11b (M1/70), CD11c (N418), CD31 (PECAM-1, 390), CD45 (30-F11), CD64, CD115 (AFS98), F4/80 (BM8), MHC II (M5/114.15.2). Biolegend, Ly6C (HK1.4) and Ly6G (1A8). Isotype-matched Abs to irrelevant epitopes were used as negative controls.

### Tissue damage markers

The activity, LDH and SDH was determined using commercial kits according to manufacturers’ instructions. Histopathology was evaluated in sections from paraffin embedded or frozen tissues using specific antibodies to nitrotyrosine as described (64).

### Quantification of hydrogen peroxide production

We measured hydrogen peroxide production using the Amplex Red Hydrogen Peroxide Kit (Invitrogen) following the manufacturer’s instructions. Neutrophils were washed and resuspended in HBSS buffer plus 1% glucose. We incubated 2×10^4^ cells with Amplex Red reaction mixture with 10^−7^ M of fMLP at 37°C for 5 min prior to measurements with a fluorimetric plate reader at an excitation wavelength of 544 nm and an emission wavelength of 590 nm. For *ex vivo* measurement of ROS production we used dihydrorhodamine 123 (65) (66). Male CD1 mice were injected with one i.p. dose of LPS [40mg/kg]; 1h and 2h later, mice were injected with fluorochrome labeled ANP or PANP as described above; 6h after LPS challenge, mice were euthanized and heart and lungs were perfused with PBS, excised lung lobes were minced and digested in collagenase solution. Erythrocytes were lysed. Leukocytes were enriched by Ficoll density gradient. Cells were resuspended in HBSS plus 1% glucose incubated with dihydrorhodamine 123 for 20 min at 37°C and then immediately processed and analyzed by flow cytometry.

### *in vivo* imaging

Surgical methods to access to the lung are based on Looney et al (28). Tail vein injection with BV421-labeled Ly6G antibody (10μg/mouse) (1A8, Biolegend) and 70kDa Tetramethylrhodamine-dextran (200μg/mice) (ThermoFisher Scientific) were performed to stain PMN and lung microvascular structures, respectively, before surgery. A resonant-scanning two photon microscope (Ultima Multiphoton Microscopes, Bruker) with an Olympus XLUMPlanFL N 20x (NA 1.00) collected dual-color images (Emission filter; 460/50 nm for Brilliant Violet 421 and 595/60 nm for Tetramethylrhodamine) with 820 nm excitation at video rate. Images were processed and analyzed by image J and customized LabVIEW programs. For PMN amount analysis, fluorescent intensities of PMN in field of view were quantified and the value of saline injected controls was normalized to 1. For PMN internalizing P/ANP, PMN number with or without P/ANP in field of view was counted and the percentage was calculated. For PMN velocity analysis, PMN velocity of cells migrating more than 1 min in the field of view, was measured.

### Adoptive transfer experiments

Donor mice, male BALB/c 8 to 10 wks, were injected with one i.p. dose of LPS [30mg/kg]. Prior to euthanasia, mice were injected, into the tail vein, with ANP containing the fluorochrome AF647. After euthanasia, both heart and lungs were perfused with PBS, excised lung lobes were minced and digested in collagenase solution. Erythrocytes were lysed. Syngeneic recipient mice of the same age, were injected, i.p., with a non-lethal dose LPS [1mg/kg] prior to adaptive transfer, i.v., of 8×10^5^ ANP^high^ or an equal number of ANP^low^ granulocytes.

### Transcriptomic profiling

Mice were injected i.p. with LPS (12 mg/kg) or a saline, ANP were injected i.v. 1h before mice were euthanized. PMN were harvested form lungs and by flow cytometry sorting of Ly6G^pos^ divided into ANP^high^ and ANP^low^ PMN. mRNA was isolated and prepared immediately for RNA-Seq or qPCR analysis. **RNA-Seq**. Raw reads were aligned to reference genome mm10 using STAR (67). Gene expression was quantified using FeatureCounts (68) against Ensemble coding and non-coding gene annotations. Differential expression between nanoparticle dye selection positive and negative was computed using edgeR (69, 70), adjusting for technical batch effect due to mouse cohort selection; normalized gene expression was reported in log2 CPM units. P-values were adjusted for multiple testing using the false discovery rate (FDR) correction of Benjamini and Hochberg (71). Significant genes were determined based on an FDR threshold of 5% (0.05). All genes that were differentially expressed due to nanoparticle dye selection, in either LPS treated or untreated animals, were visualized in a heatmap, including dendrograms from complete linkage hierarchical clustering for both genes and samples. In addition, separate heatmaps for chemokine ligands and chemokine receptors were generated, plotting all genes with CPM > 0.25 (10 reads at a sequencing depth of 40M reads) regardless of differential expression levels. LPS treated animals were separated from untreated animals in these heatmaps to highlight the effect of nanoparticle dye selection on the expression levels. **qPCR**. mRNA was extracted from sorted PMN using the Qiagen RNeasy Mini Kit. Total RNA quantity was measured at 260nm and purity was assessed by the optical density 260nm/optical density 280nm ratio. 0.5μg of RNA was transcribed to complementary DNA with random primers using the High-Capacity cDNA Reverse Transcription Kit (ThermoFischer). Quantitative gene expression was evaluated by qPCR using the QuantStudio 7 Flex Real-Time PCR System. Results were calculated using the comparative C_T_-method(72), and expressed relative to the expression of the housekeeping gene Ppia (ENSMUSG00000071866.12). We used the following primers, forward, and reverse, respectively: Ppia GGCAAATGCTGGACCAAACAC, TTCCTGGACCCAAAACGCTC: Il1b TGGGAAACAACAGTGGTCAG, CAAGGAGGAAAACACAGGCT; Ccr1 CAACCTGGCTGTCTCTGATC, GCATGGAAGCTAAGATGGCT; Il15 CAATTCTCTGCGCCCAAAAG, TCTTAAGGACCTCACCAGC; Ccl3 AGAAGGATACAAGCAGCAGC, GACTTGGTTGCAGAGTGTCA; Ccl4 GATCTGTGCTAACCCCAGTG, AGAAGAGGGGCAGGAAATCT; Cxcl2 AGTTTGCCTTGACCCTGAAG, GTGAACTCTCAGACAGCGA; Cxcl3 GCCCCAGGCTTCAGATAATC, AAAGACACATCCAGACACCG.

### Statistical analysis

We examined the differences between groups for statistical significance by Student’s t-test or ANOVA, and compared survival curves with a log-rank test. Enrichment of chemokine receptor or chemokine expression in the ANP^high^ and ANP^low^ groups was assessed by Fisher’s Exact Test. A *p* value of <0.05 was considered statistically significant.

## Acknowledgements

We thank Dr. Paul Frenette of Albert Einstein College of Medicine for helpful comments.

## Video

Two-photon microscopy of perfused, ventilated lungs from endotoxemic mice and saline-injected controls. mice were injected with LPS [30mg/kg], 6h prior to, and with ANP or PANP 3h and 4h prior to *in vivo* imaging. Dextran traces blood vessels, outlining lung microvascular structures, blue; fluorescent-labeled Ly6G Abs label PMN, green; fluorochrome AF647 labels ANP or PANP, red.

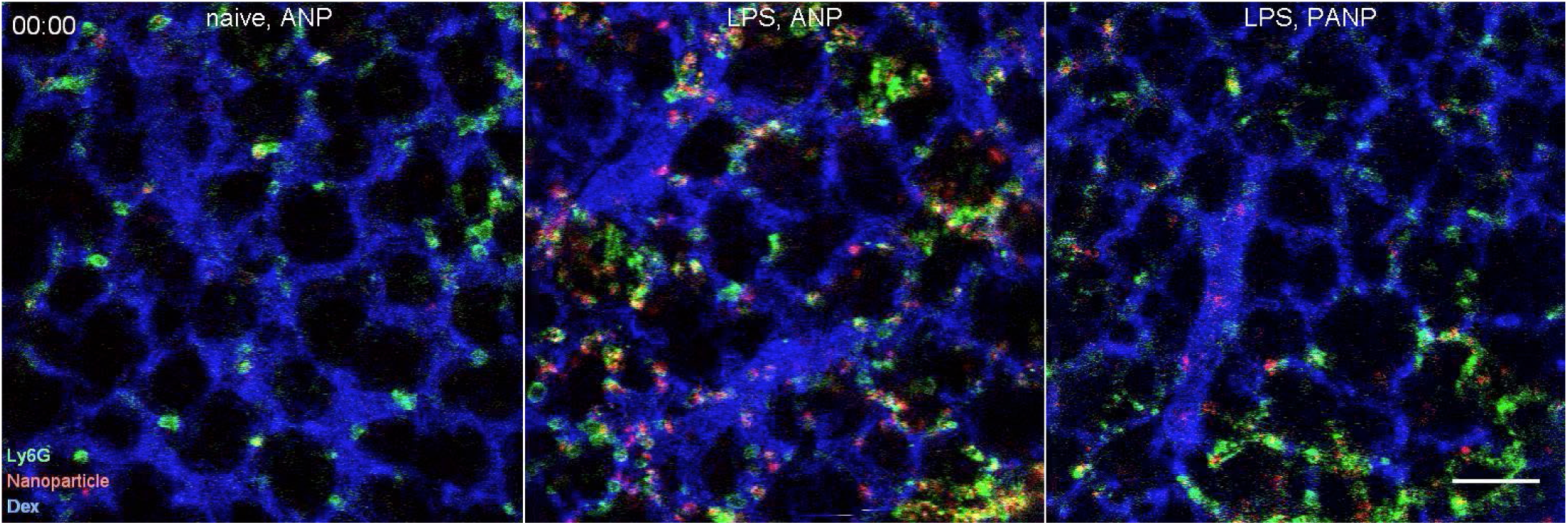

Colors are pseudocolors. Bar measures 50μm. Representative data from a minimum or 4 mice per treatment group.

## Supplemental Figures

**Supplemental Figure 1.**
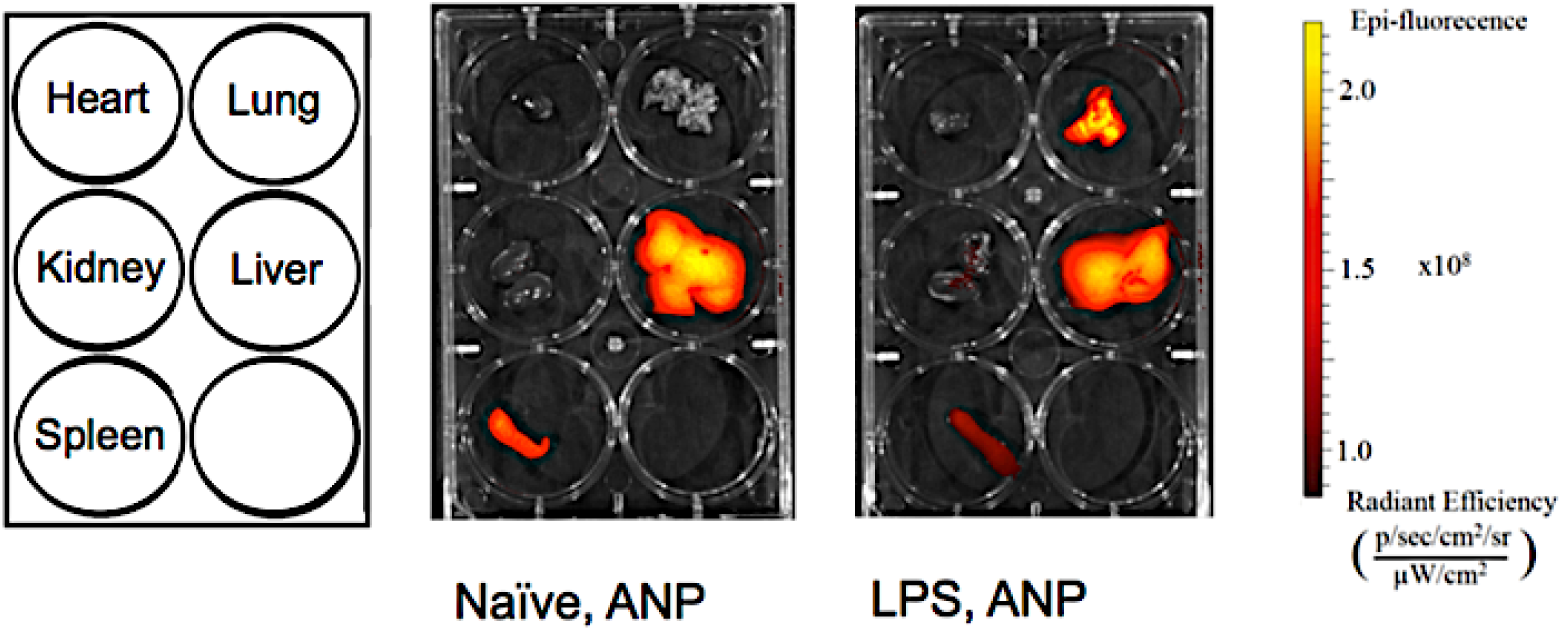
ANP Biodistribution Measured by IVIS. In naïve mice, i.v.-injected ANPs were mostly found to liver and spleen. After i.p. administration of the endotoxin LPS [30mg/kg], ANP appeared prominently in lungs, and remained visible in liver and in spleen. Albumin was labeled with vivotag 800 (PerkinElmer) and then formed into ANPs. Organs were harvested 4 h after 150µl tail vein injection of 2 mg/ml vivotag 800 labeled ANPs. Epi-fluorescence radiance scale corresponding to images. Fluorescence was measured by a Xenogen IVIS Spectrum (Caliper Life Sciences) and images were processed with Living Image software (ver. 4.3.1). An excitation filter of 785nm and emission filter of 820nm with 120 sec exposure times were used for all experiments. Representative data obtained from at least 3 mice per treatment group.

**Supplemental Figure 2.**
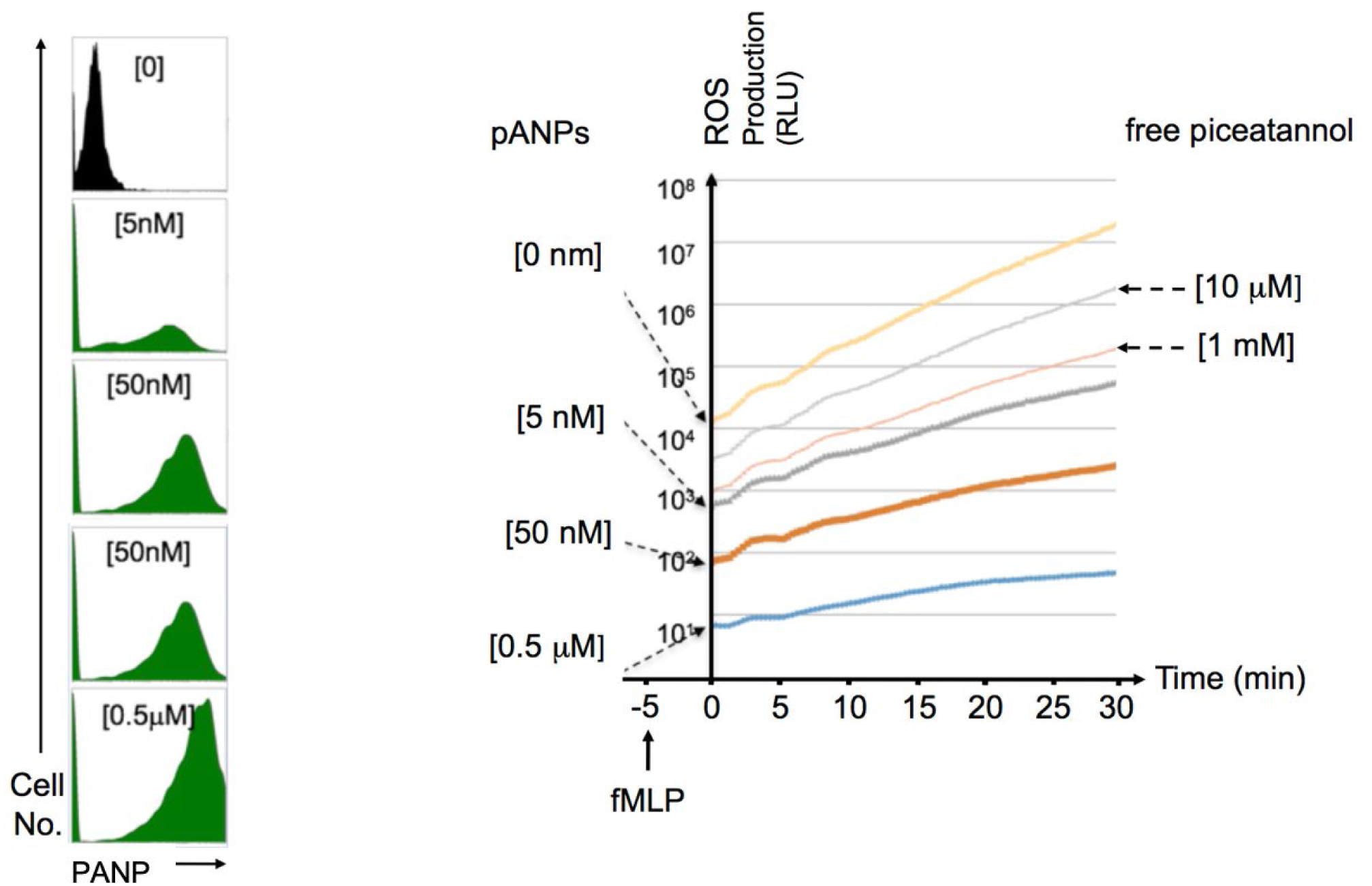
**Endocytosis of PANP (left panel).** Flow cytometry histograms of bone marrow PMN stimulated with LPS [1mg/ml] and incubated with PANP, at the indicated doses, for 30 min at 37°C, and then processed for flow cytometry. Representative data obtained from at least 3 mice per treatment group **PANP treatment reduces superoxide production by PMN in response to fMLP stimulation.** PAMP reduced fMLP stimulated ROS production in a dose dependent manner. Bone marrow PMN were incubated with the indicated dises of PANP and stimulated with fMLP for the indicated time. ROS production was measured using an Amplex Red Hydrogen Peroxide/Peroxidase Assay Kit. Representative data obtained from at least 3 mice per treatment group.

**Supplemental Figure 3.**
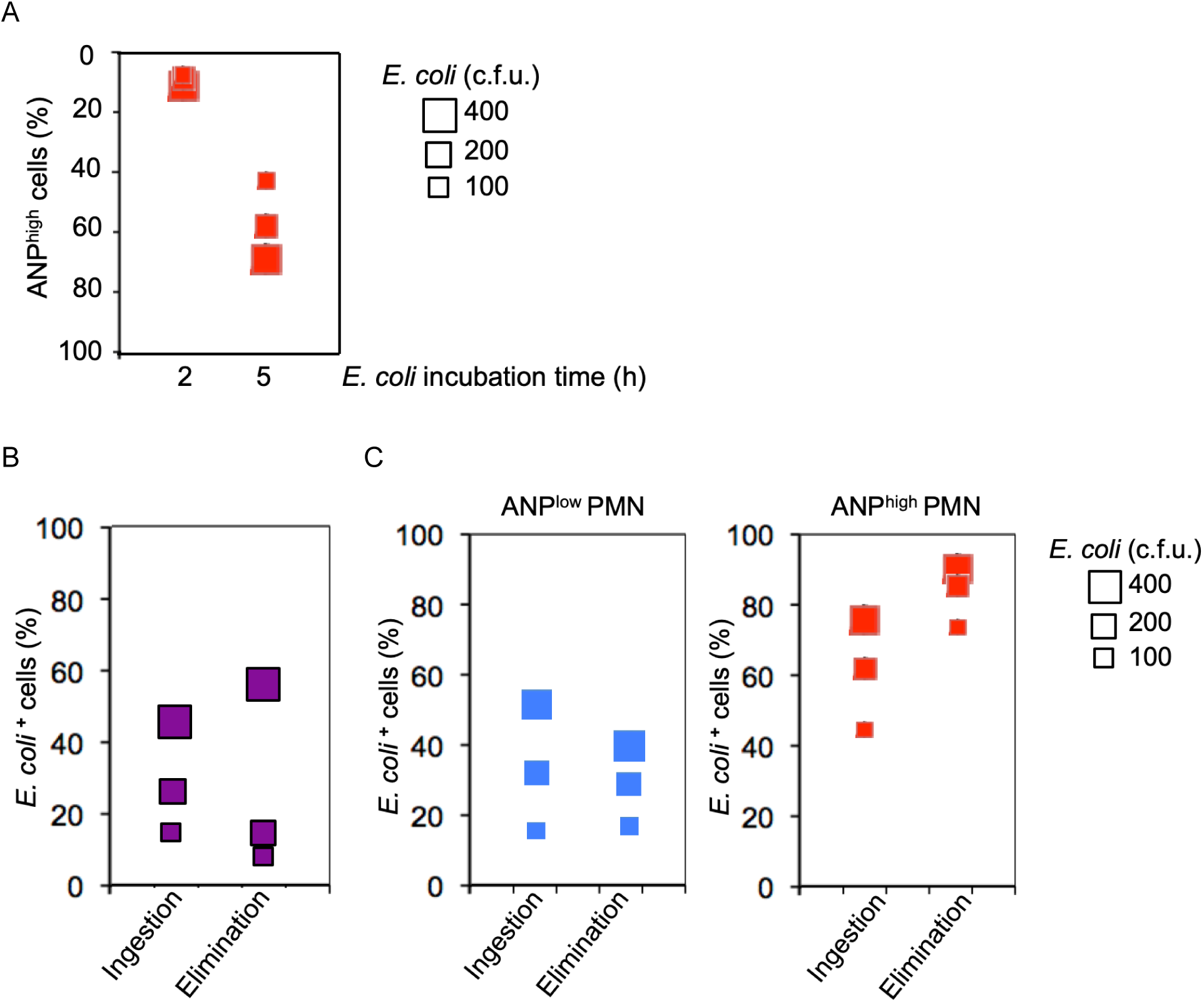
Neutrophil heterogeneity determines endocytic functions. (**A**) **Live *E. coli* bacteria prime Ly6G**^**+**^ **bone marrow neutrophils for endocytosis of albumin nanoparticles (ANP)**. Endocytosis, expressed as percentage of ANP^+^ cells, increased with the time of exposure and with the dose of bacteria. Endocytosis assay. Albumin nanoparticles (ANP) labeled with AF647 were incubated with 1×10^6^ bone marrow cells for 15min at 37°C at the end of their priming with increasing numbers of viable (c.f.u.) DH5α *Escherichia coli* (*E. coli*) bacteria. After incubation, cells were fixed and prepared for flow cytometric analysis. (**B**) **Elimination of live *E. coli* bacteria is dose-dependent.** Challenge with the highest number of live *E. coli* bacteria overwhelmed elimination of bacteria by Ly6G^+^ bone marrow neutrophils. Ingestion, percentage of GFP^+^ cells after 2h of incubation with *E. coli* bacteria expressing GFP. Elimination, percentage of GFP^+^ cells after 5h of incubation with *E. coli* bacteria expressing GFP. **(C) Heterogeneous response of neutrophils to *E. coli* challenge.** The subset of neutrophils that is inefficient in eliminating *E. coli* bacteria is susceptible to ANP uptake (ANP^high^). Neutrophils resistant to ANP-uptake (ANP^low^) are efficient in elimination of *E. coli* bacteria. Phagocytosis assay. Increasing numbers of viable (c.f.u.) DH5α *Escherichia coli* (*E. coli*) bacteria expressing GFP were incubated with 1×10^6^ bone marrow cells for 2h or 5h at 37°C. After incubation, cells were fixed and prepared for flow cytometric analysis. Specific Ab to neutrophil-marker Ly6G was used.

## Supplemental Table

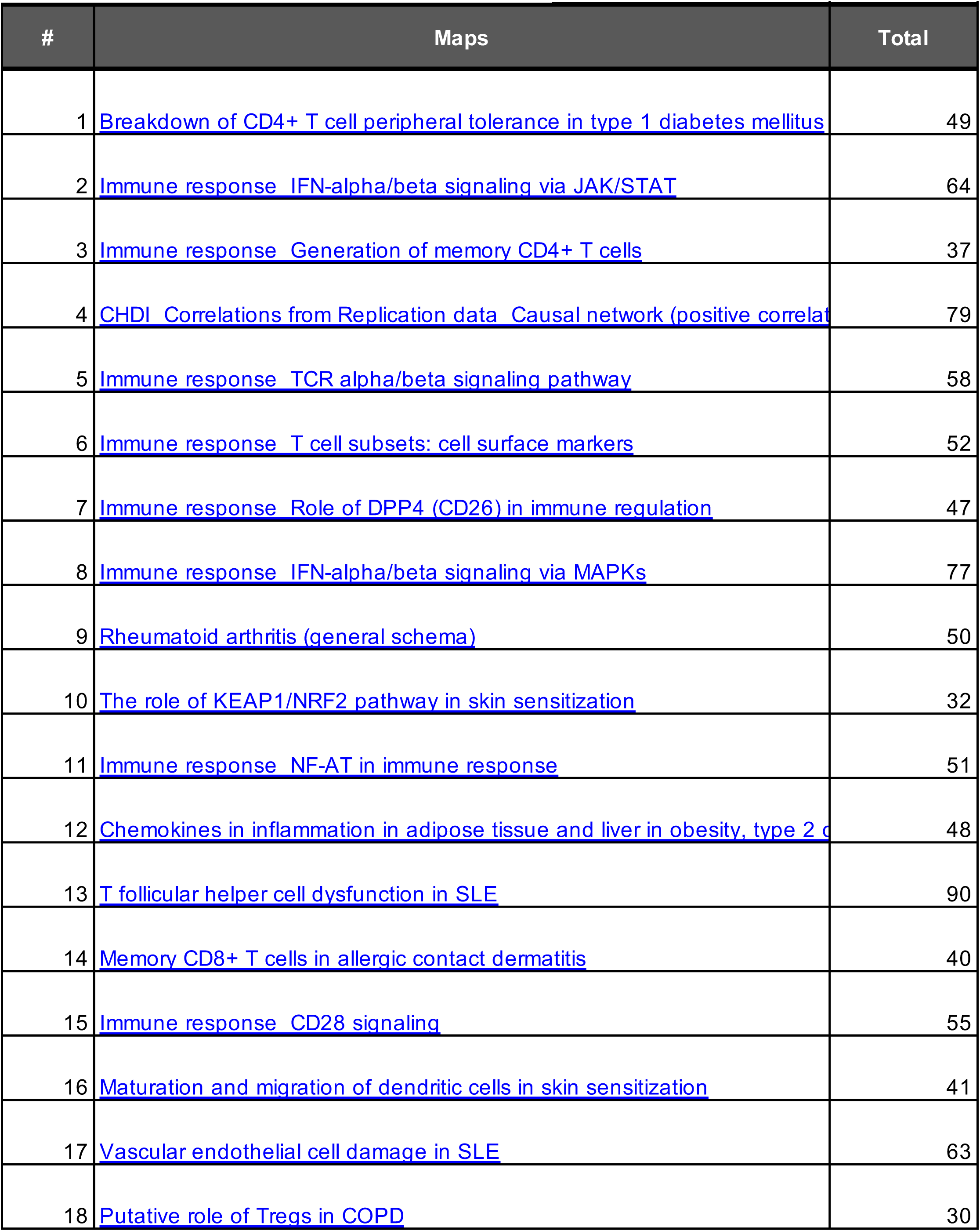
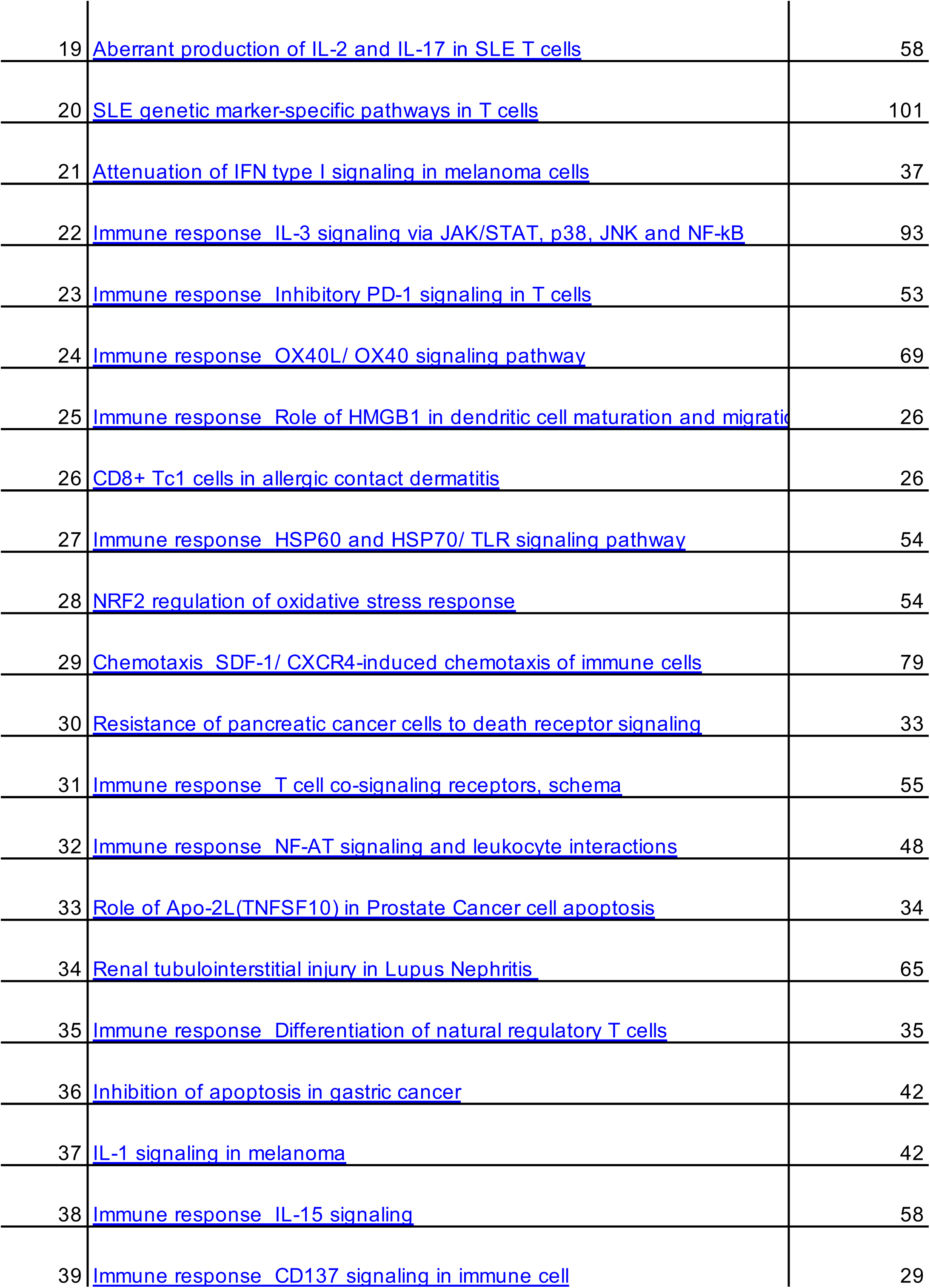
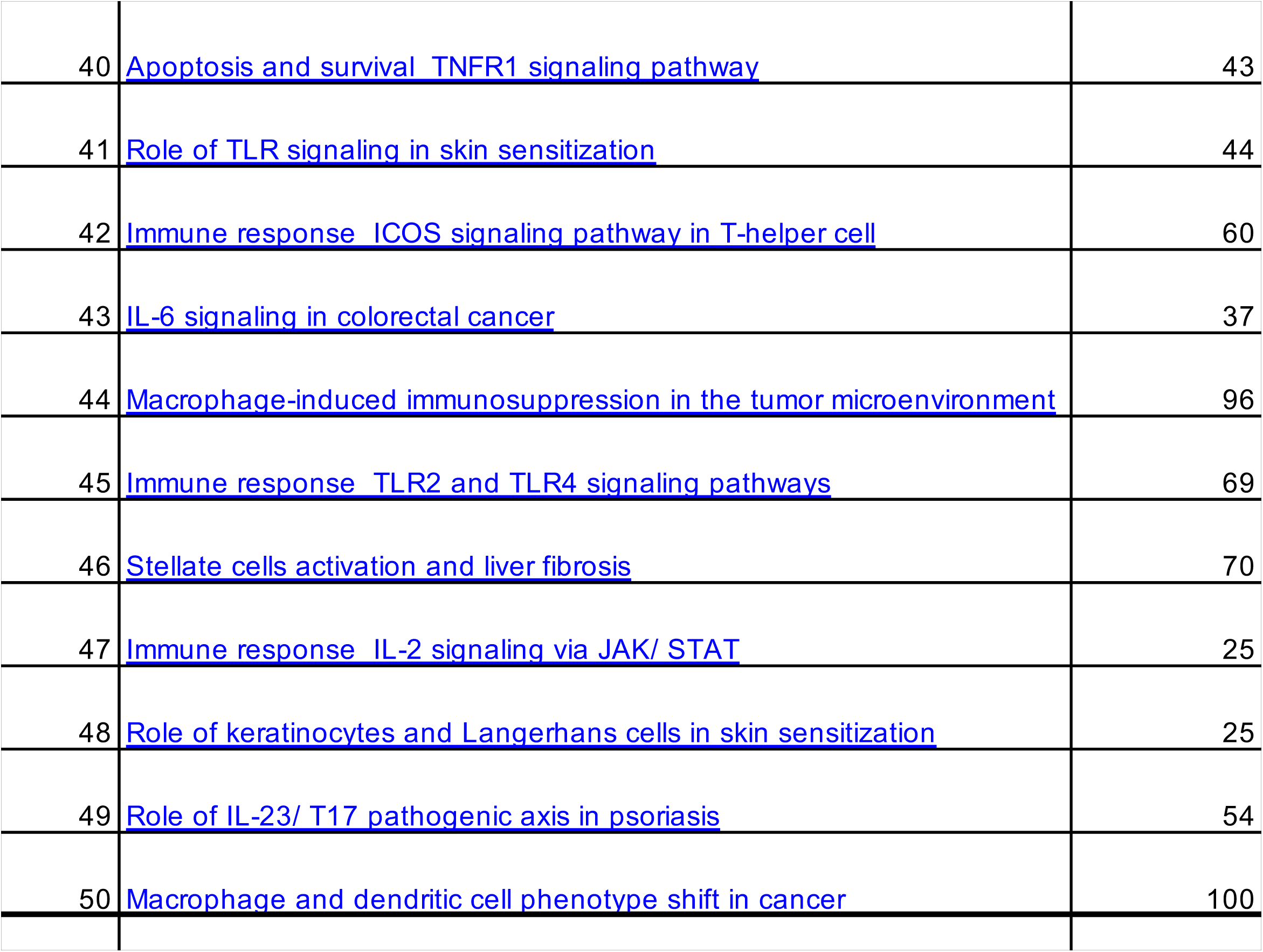
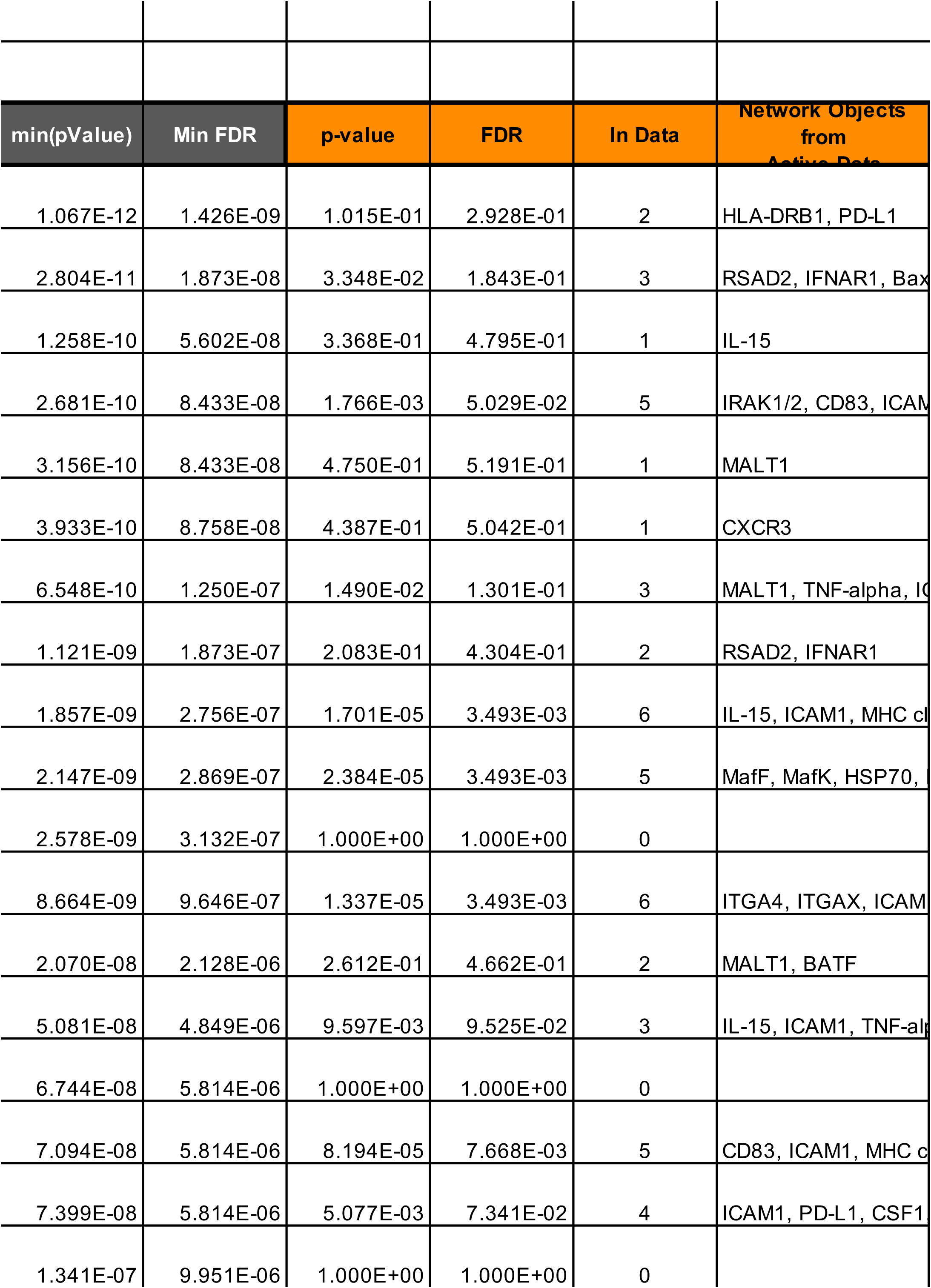
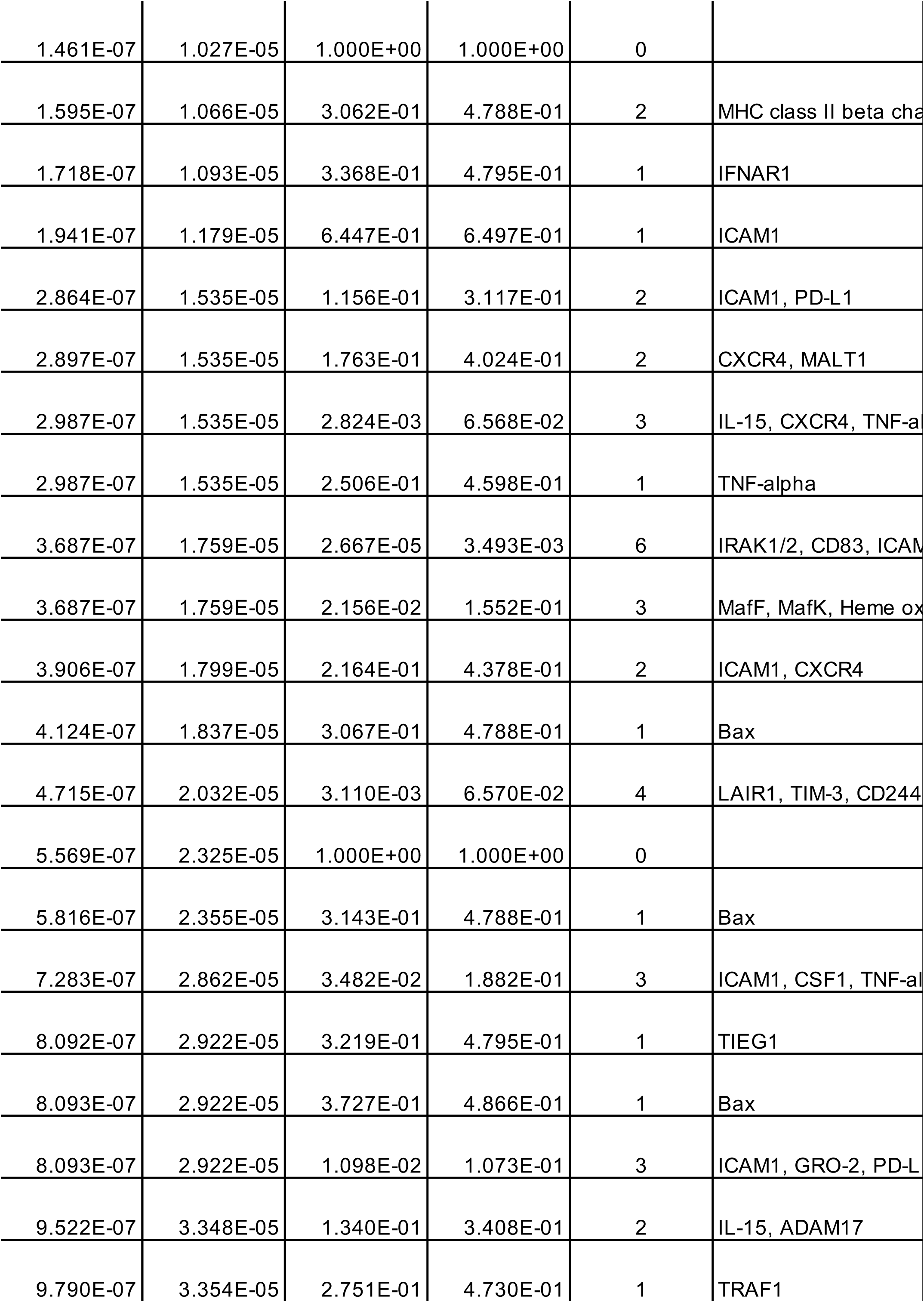
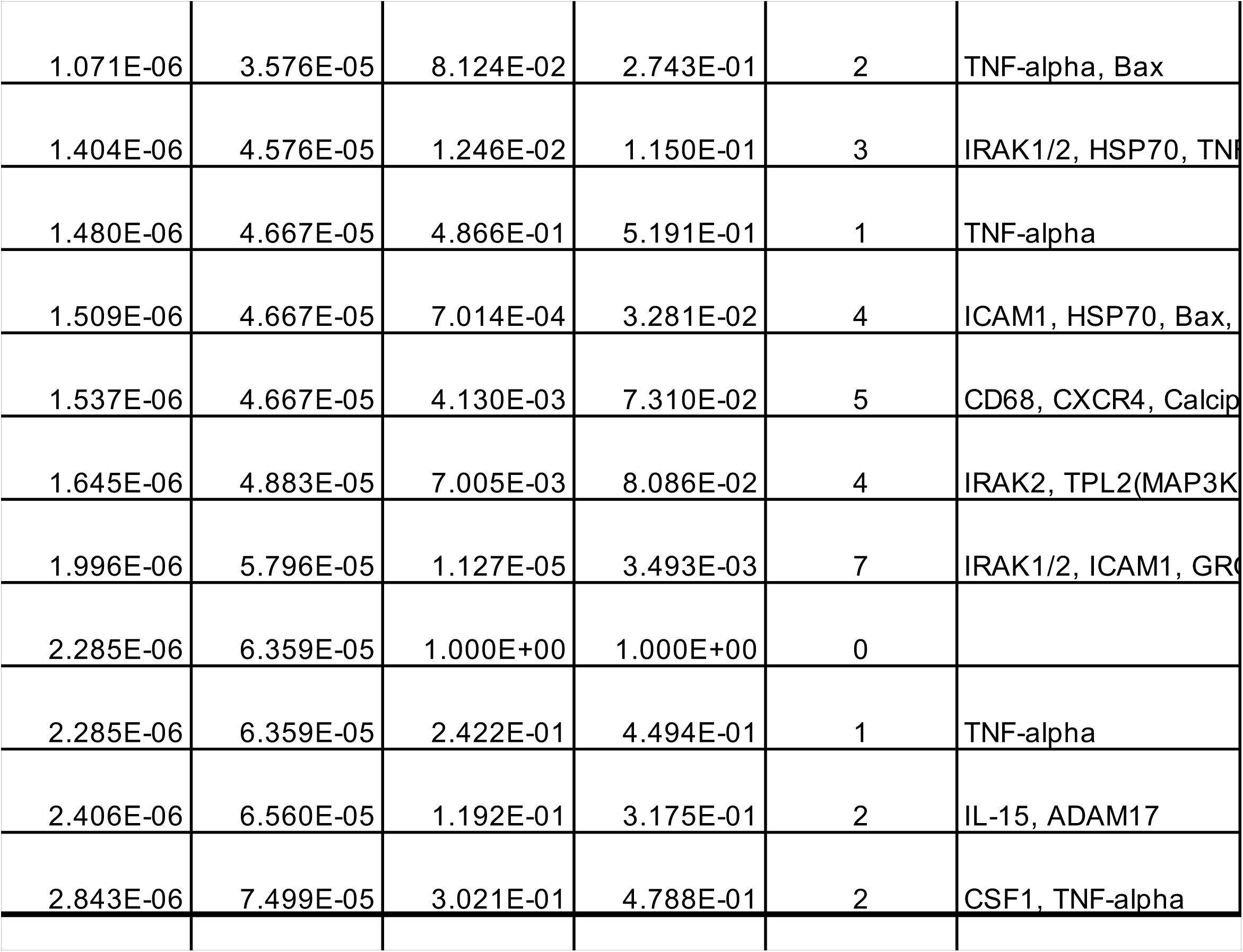
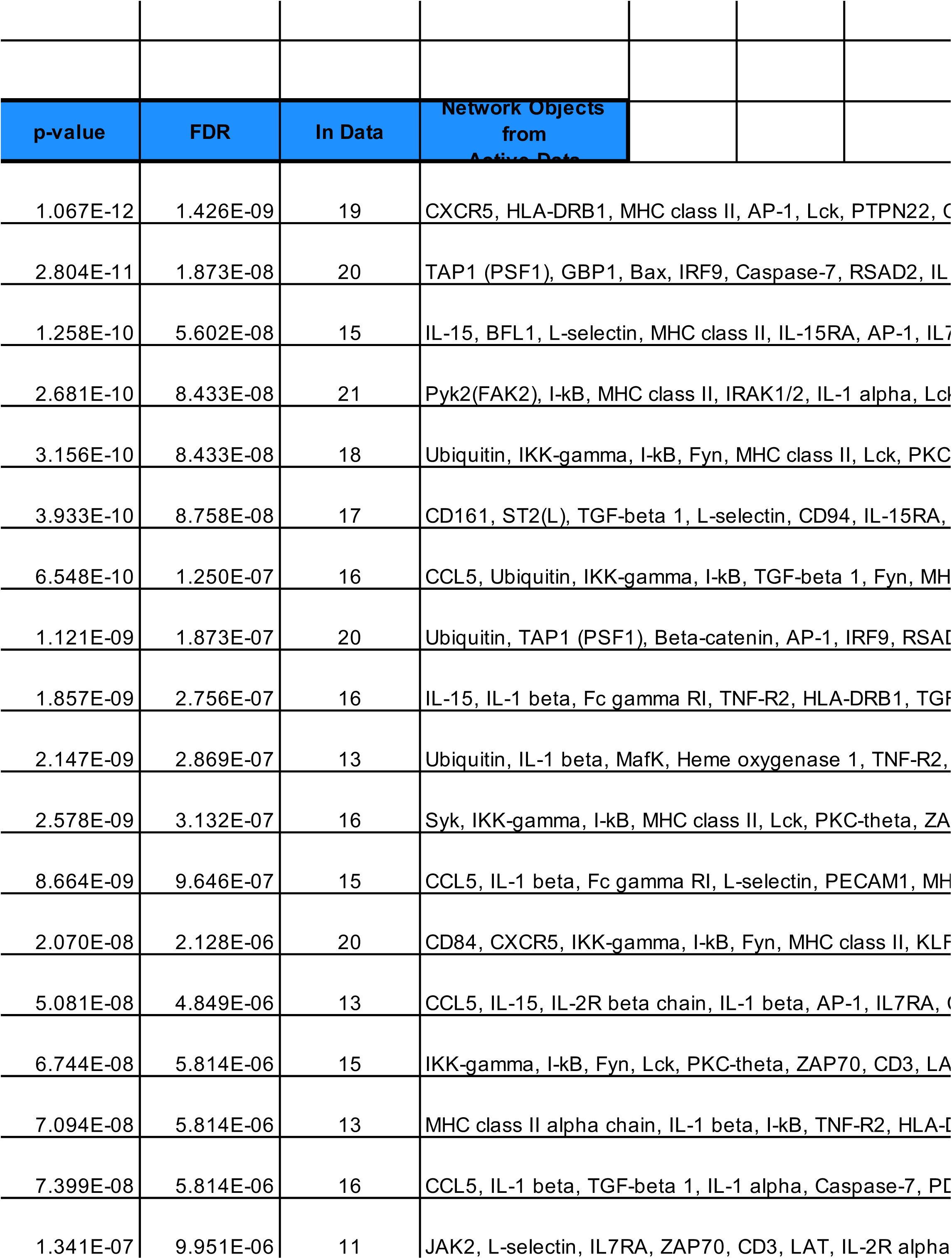
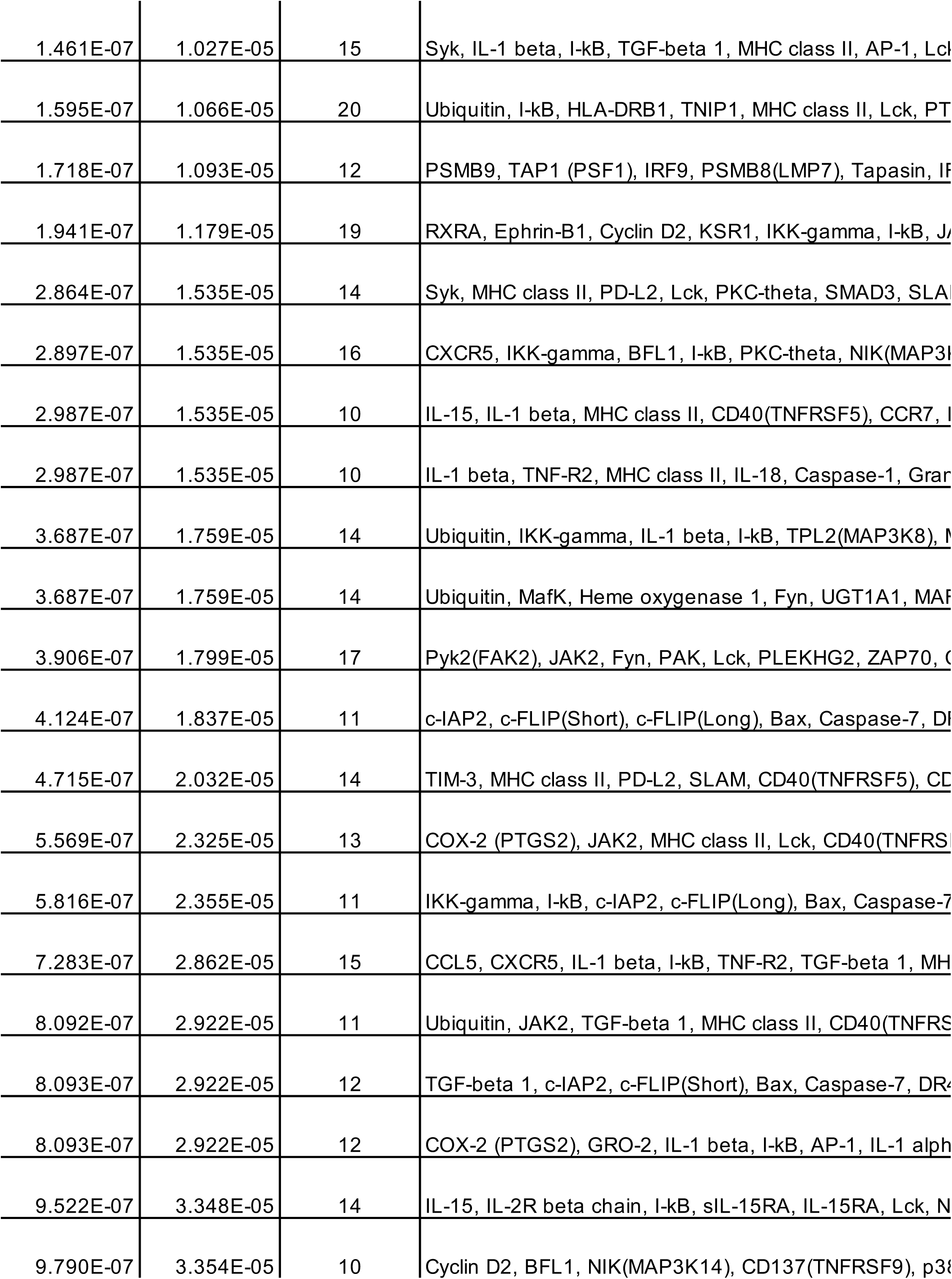
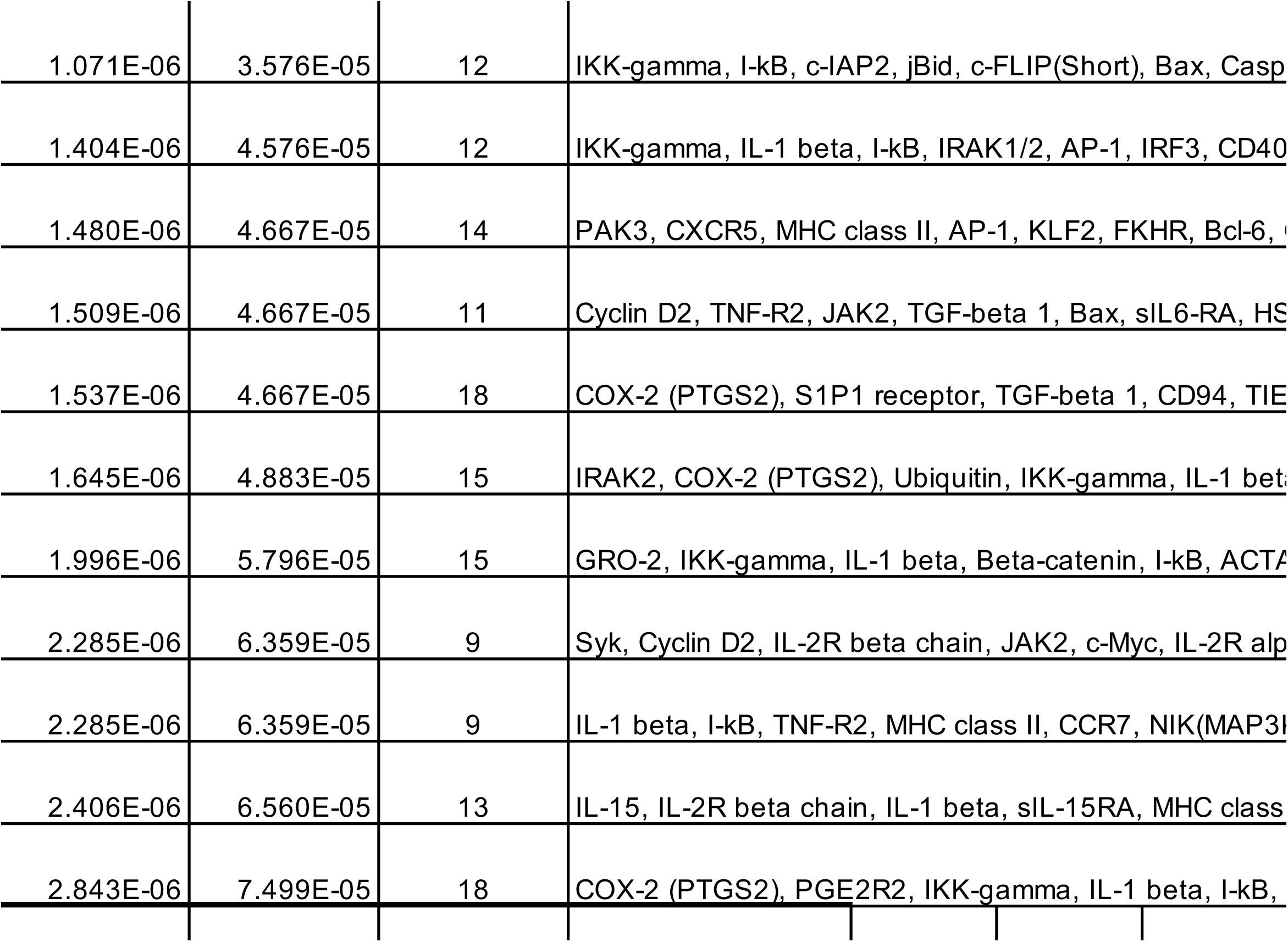
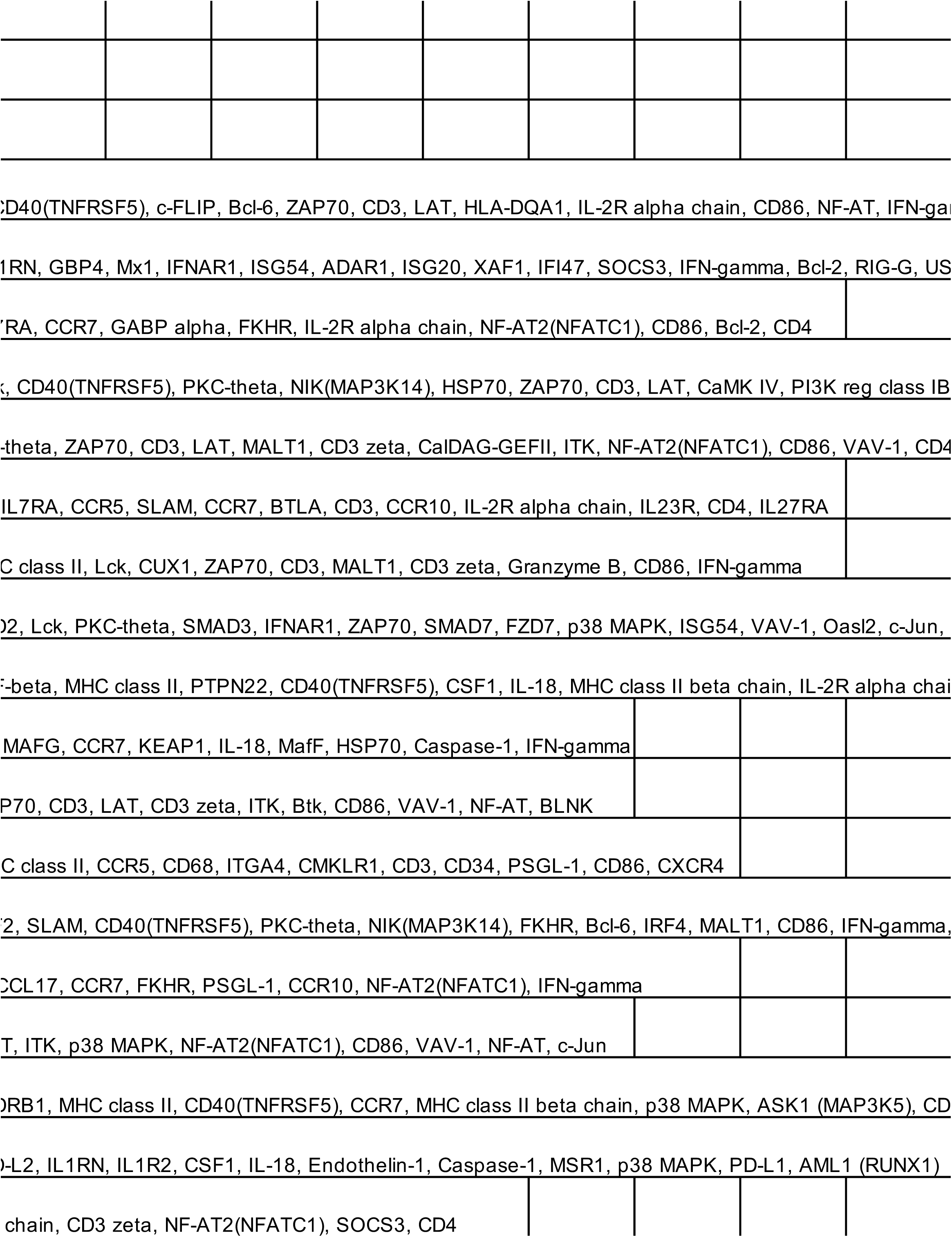
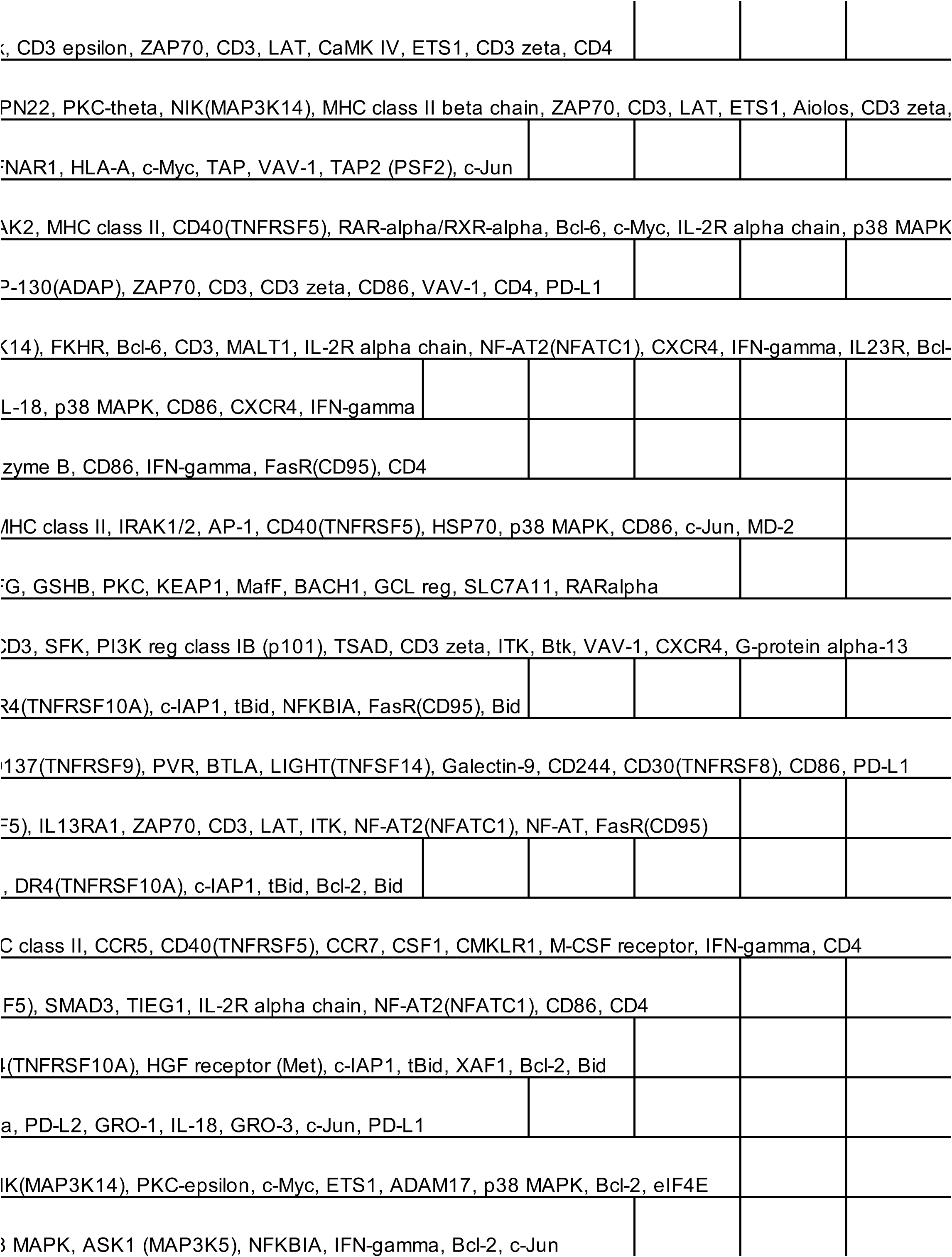
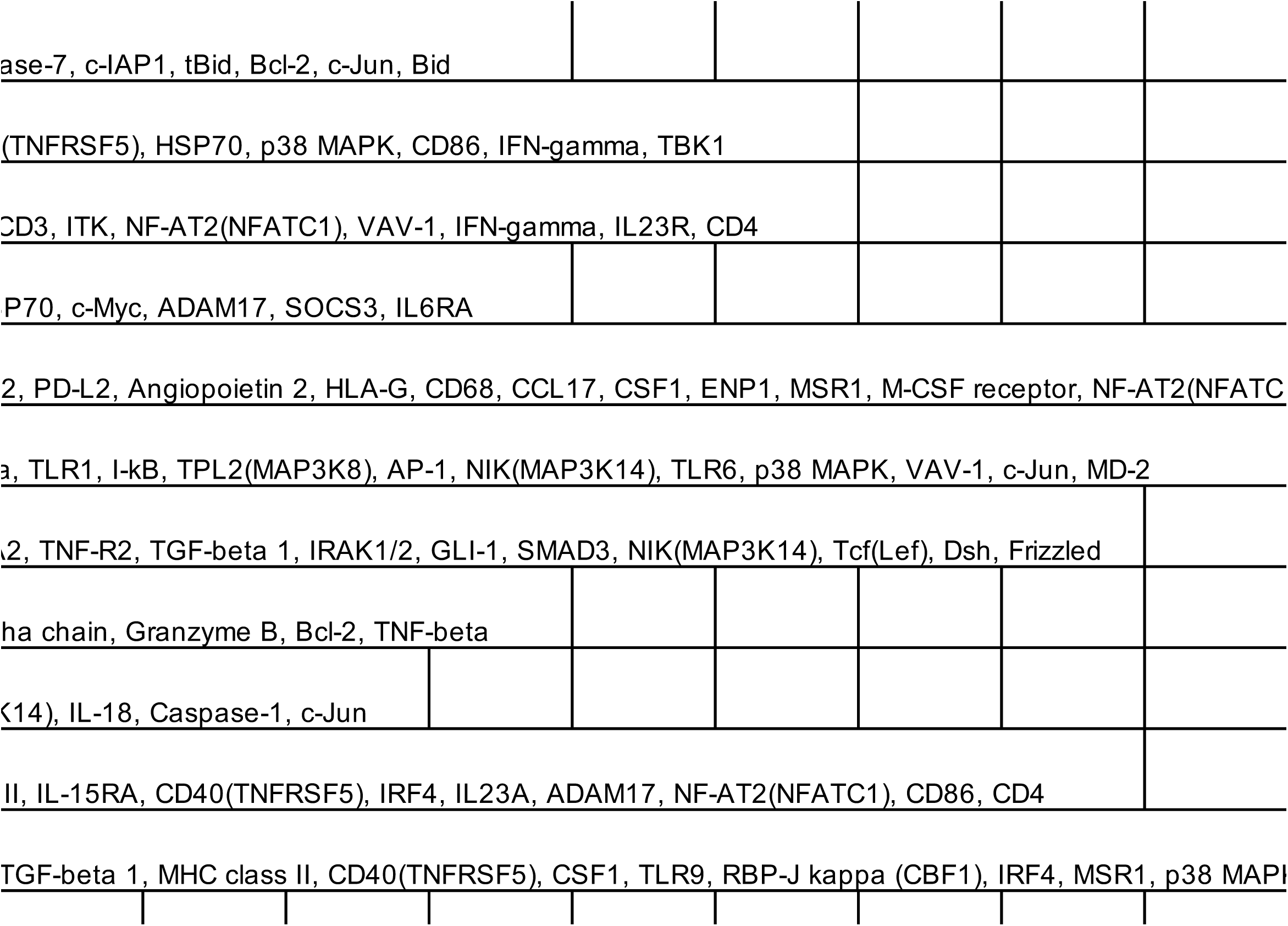
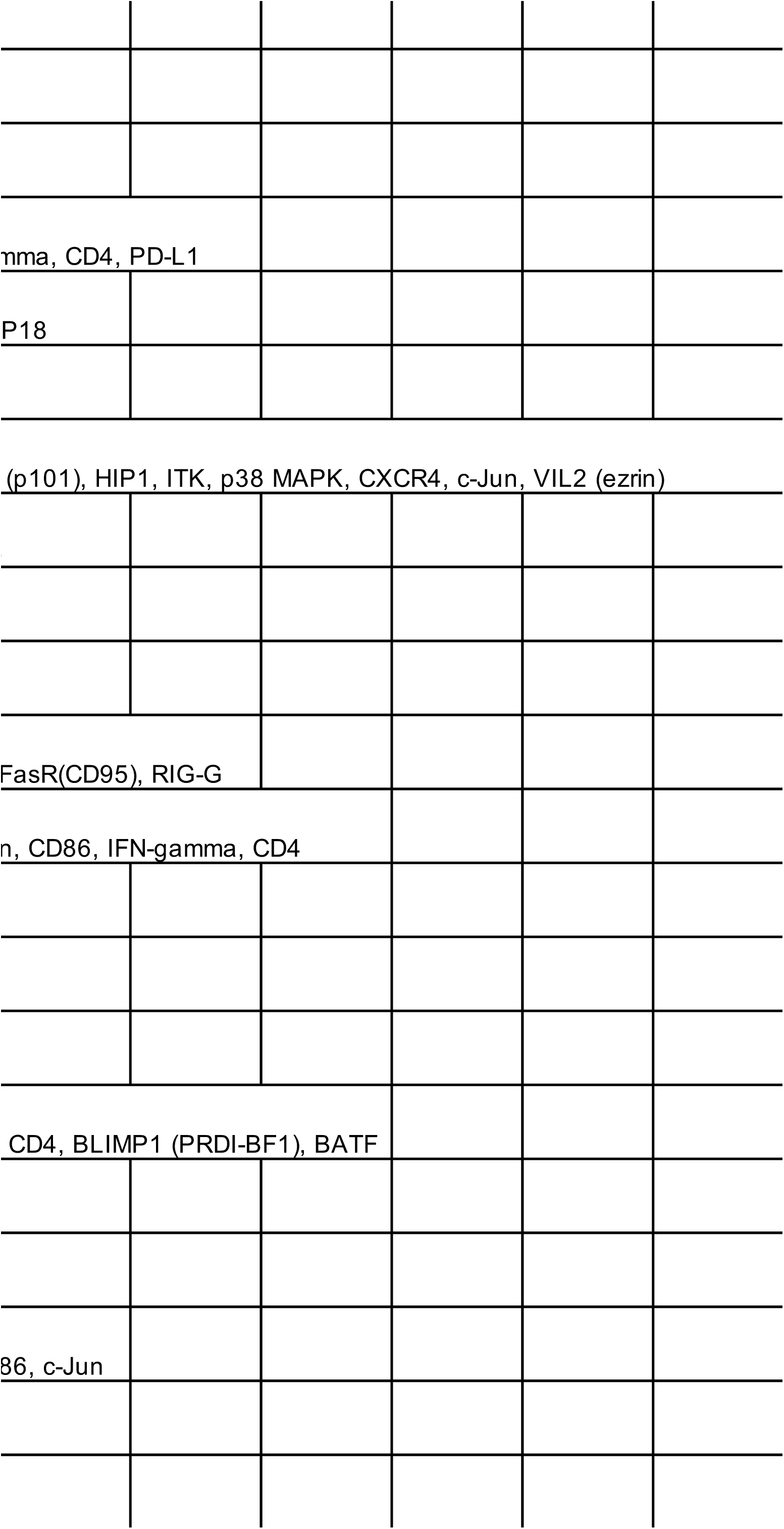
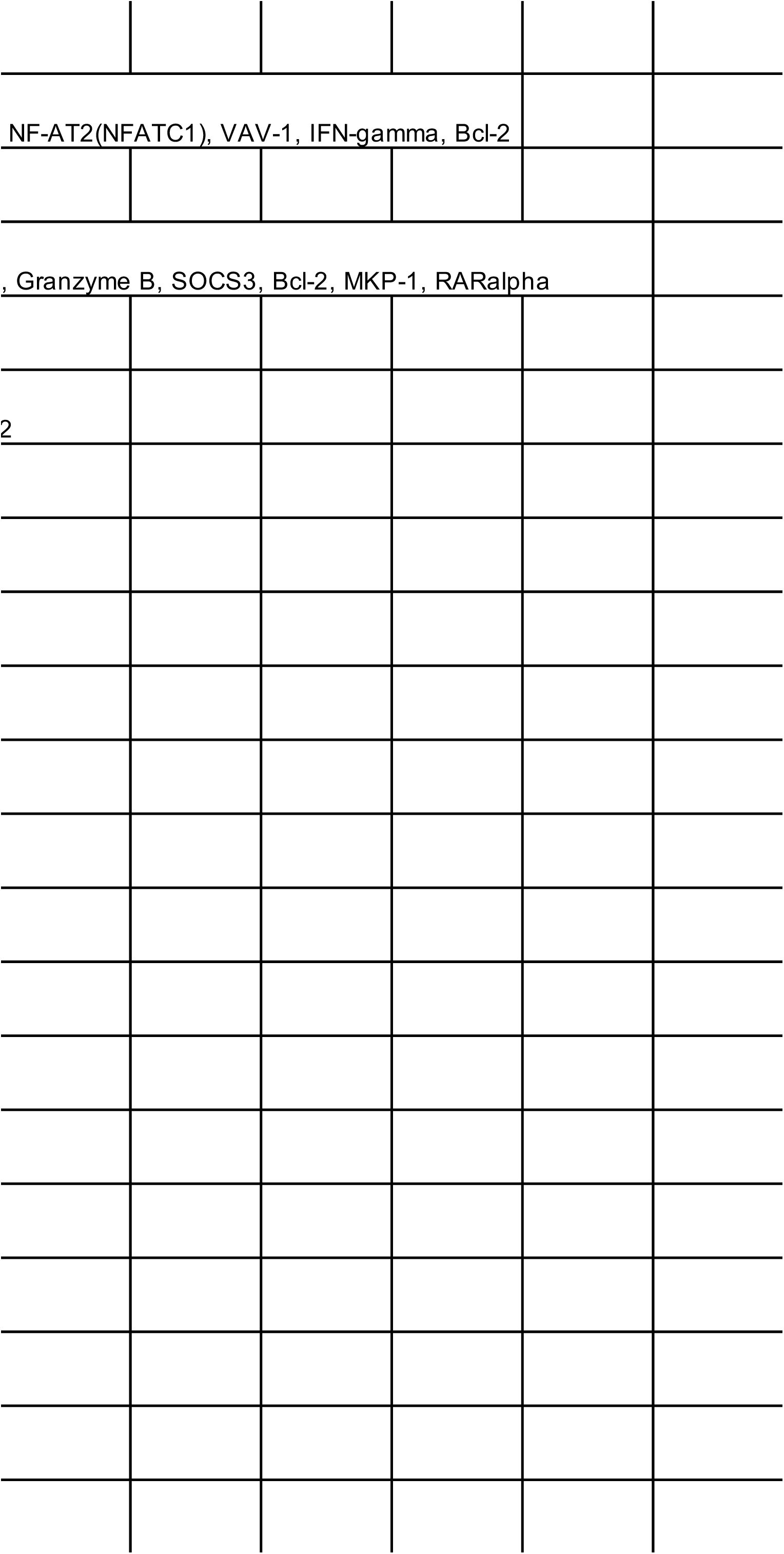
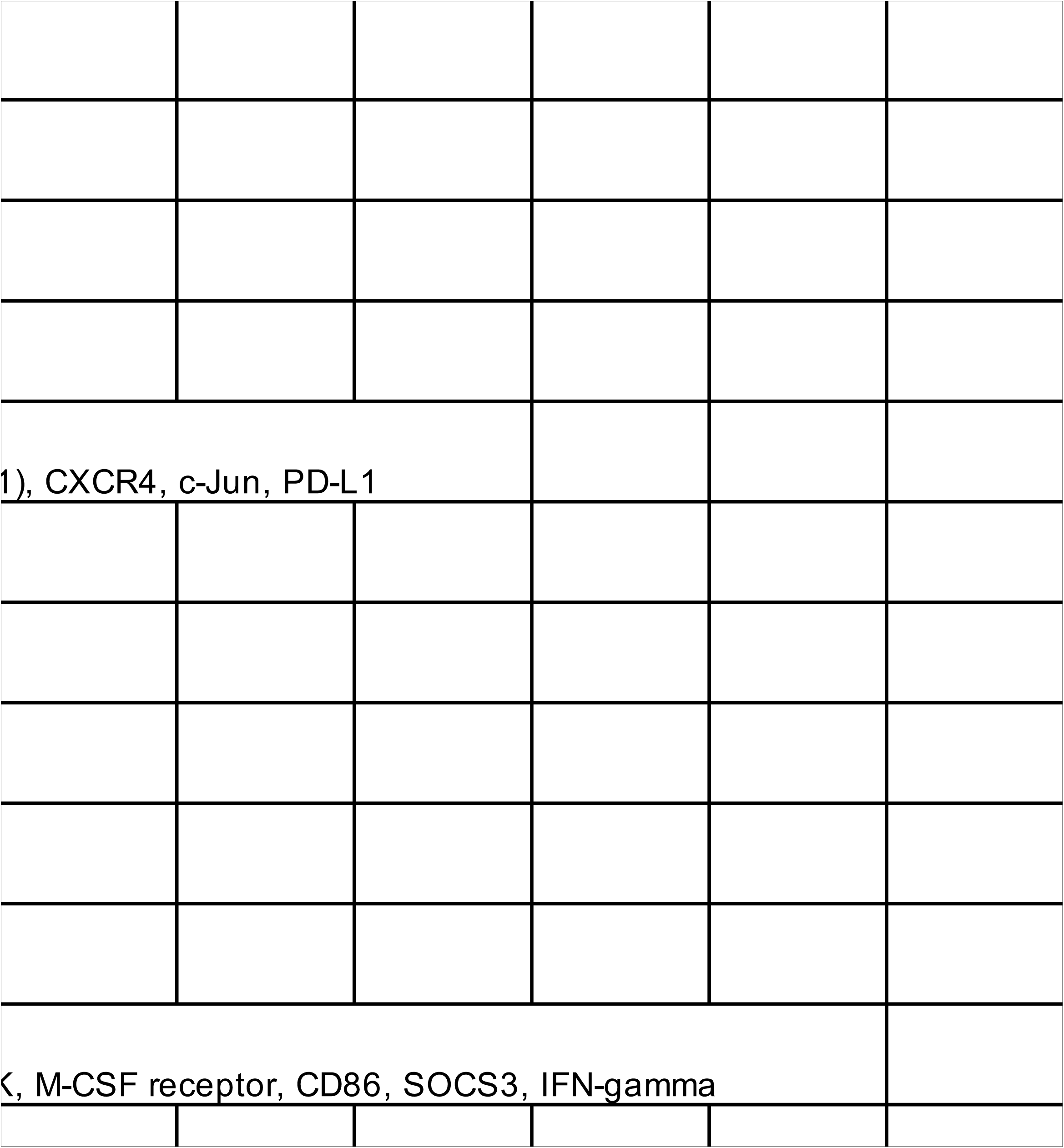

